# Comparative transcriptomics in ferns reveals key innovations and divergent evolution of the secondary cell walls

**DOI:** 10.1101/2024.08.27.609851

**Authors:** Zahin Mohd Ali, Qiao Wen Tan, Peng Ken Lim, Hengchi Chen, Lukas Pfeifer, Irene Julca, Jia Min Lee, Birgit Classen, Sophie de Vries, Jan de Vries, Teng Seah Koh, Li Li Chin, Fanny Vinter, Camille Alvarado, Amandine Layens, Eshchar Mizrachi, Mohammed Saddik Motawie, Bodil Joergensen, Peter Ulvskov, Yves van de Peer, Boon Chuan Ho, Richard Sibout, Marek Mutwil

**Affiliations:** School of Biological Sciences, Nanyang Technological University, Singapore, Singapore; Department of Plant Biotechnology and Bioinformatics, Ghent University, 9052 Ghent, Belgium; VIB Center for Plant Systems Biology, VIB, 9052 Ghent, Belgium; Pharmaceutical Institute, Department of Pharmaceutical Biology, Christian-Albrechts-University of Kiel, Kiel, Germany; University of Lausanne (UNIL), Genopode 2024.3, 1015 Lausanne, Switzerland; Department of Applied Bioinformatics, Institute for Microbiology and Genetics, Goettingen Center for Molecular Biosciences (GZMB), Campus Institute Data Science (CIDAS), University of Goettingen, Goldschmidtstrasse 1, D-37077, Göttingen, Germany; Singapore Botanic Gardens, National Parks Board, 1 Cluny Road, Singapore, 259569, Republic of Singapore; INRAE, UR BIA, F-44316 Nantes, France; Department of Biochemistry, Genetics and Microbiology, Forestry and Agricultural Biotechnology Institute (FABI), University of Pretoria, Pretoria 0002, South Africa; Department of Plant and Environmental Sciences, University of Copenhagen, Frederiksberg, Denmark; College of Horticulture, Academy for Advanced Interdisciplinary Studies, Nanjing Agricultural University, Nanjing, China; Department of Biological Sciences, National University of Singapore, Republic of Singapore

## Abstract

Despite ferns being crucial to understanding plant evolution, their large and complex genomes has kept their genetic landscape largely uncharted, with only a handful of genomes sequenced and sparse transcriptomic data. Addressing this gap, we generated extensive RNA-sequencing data for multiple organs across 22 representative species over the fern phylogeny, assembling high-quality transcriptomes. These data facilitated the construction of a time-calibrated fern phylogeny covering all major clades, revealing numerous whole-genome duplications and highlighting the uniqueness of fern genetics, with half of the uncovered gene families being fern-specific. Our investigation into fern cell walls through biochemical and immunological analyses identified occurrences of the lignin syringyl unit and its independent evolution in ferns. Moreover, the discovery of an unusual sugar in fern cell walls hints at a divergent evolutionary path in cell wall biochemistry, potentially driven by gene duplication and sub-functionalization. We provide an online database preloaded with genomic and transcriptomic data for ferns and other land plants, which we used to identify an independent evolution of lignocellulosic gene modules in ferns. Our data provide a framework for the unique evolutionary path that ferns have navigated since they split from the last common ancestor of euphyllophytes more than 360 million years ago.

## Introduction

Since they diverged from a shared ancestor with seed plants more than 360 million years ago, ferns have played a significant role in life on Earth ^1^. They occupy various niches in different ecosystems, acting as pioneer species, key ecological players, invasive entities, and contributors to agriculture. They are the second most diverse group of vascular plants after angiosperms, with over 10,500 existing species ^2–6^. Ferns exhibit great morphological and physiological diversity, and have evolved equally diverse strategies to cope with environmental challenges ^7^, such as adaptations to low-light environments ^8^. The secondary metabolites produced by ferns and the genes responsible for their biosynthesis are of great interest for environmental clean-up efforts, agriculture, and the discovery of new pharmaceuticals ^9–11^.

Despite fern’s ecological importance, the phylogenetic relationship of major clades in Monilophyta (ferns) remains elusive ^12^. Based on a Maximum Likelihood (ML) tree of a concatenated matrix of 146 low-copy nuclear genes, Qi and coauthors ^13^ inferred Marattiales to be sister to Polypodiidae (i.e. the leptosporangiate ferns) as proposed in Pteridophyte Phylogeny Group (PPG) I (2016). Conversely, Shen and coauthors ^12^ inferred Marattiales to be sister to Ophioglossidae (consisting of Psilotales and Ophioglossales), based on a coalescent-based tree of two low-copy nuclear gene sets of 69 transcriptomes. Nitta and coauthors ^14^ inferred Gleicheniales as a monophyletic clade based on a ML tree of a concatenated matrix of 79 plastome loci as opposed to the paraphyletic inference of Shen and coauthors ^12^ and Qi and coauthors ^13^. It is also unclear whether horsetails are a sister group to the last common ancestor of all the remaining ferns or other fern clades, such as Marattiales ^5,12^.

Given ferns’ critical evolutionary position as the sister group to seed plants, investigating their genomes, coding sequences, and gene families offers unparalleled insights into the evolution of plants ^15^, especially the key aspects of vasculature and cell walls. The evolution of vasculature and secondary cell walls precipitated a 10-fold increase in plant species numbers (http://www.theplantlist.org/) and shaped the Earth’s geo- and biosphere ^16^. Ferns thus harbour key information for the evolution of vascular plant form and function ^17^.

However, ferns are infamous for their exceptionally large genomes (on average 12.3 billion base pairs), with one of the largest genome of any living organism - 160 billion base pairs - found in ferns^18^. They also have exceptionally high numbers of chromosomes (averaging at 40.5, with a peak at 720)^19^, which are believed to result from multiple instances of whole-genome duplication ^20–22^ and a relatively slow genome downsizing process ^23^. Among plants, ferns show the highest rate of polyploidy-driven speciation ^24^, a direct relationship between genome size, chromosome number and the age of long terminal repeat-retrotransposon (LTR-RT) insertions ^25–27^, and a high rate of whole genome duplications (WGDs) among several fern lineages ^23,28^. However, the genetic and genomic evidence for widespread whole-genome duplication in ferns remains largely unexplored ^29–31^.

Thus, comparative studies that investigate the evolution of ferns have been hampered by their large, complex genomes, limiting our understanding of fern genome evolution and the genetic underpinnings of the evolution of vasculature and cell walls. To date, only few fern genomes and transcriptomes are available ^20,32–35^, and no studies that conducted a comprehensive comparison of gene inventories, transcriptional programs and biochemical properties of their cell walls have been reported.

To address this, we generated 405 RNA-sequencing samples to generate coding sequences and gene expression atlases for 22 fern species, capturing major representatives of ten fern orders. We investigated ancient polyploidy, the distribution of fern-specific gene families, how gene age correlates with organ-specific expression, and predicted the functions of fern-specific genes. To better understand how fern cell walls have evolved, we performed a comprehensive histological and biochemical analysis of fern tissues and propose a novel biosynthetic pathway of lignocellulose. We further detected a wide-spread occurrence of the unusual hemicellulose mixed-linkage glucan, and show that it evolved independently in ferns, and propose candidate glucosyltransferases responsible for its synthesis. We also detected a novel type of a methylated sugar, a 2-O-Methyl-D-glucopyranose. We also show that ferns likely independently evolved secondary cell walls through several duplication events in the cellulose synthase family. Finally, we make our fern genomic and transcriptomic data easily accessible with the CoNekt database (https://conekt.plant.tools/).

Our data, findings, and tools shed light on the evolution of cell walls, lignin, specialised metabolism, and organ development in ferns and other land plants. We envision that similar large, comparative studies will elucidate the evolution of plants and other organisms.

## Results

### Construction of fern coding sequences by transcriptome assembly

To capture the diversity of ferns, we selected 22 candidate species representing ten orders, which were photographed and dissected on site (Figure S1), with organs categorised with localities and vouchers attached (Table S1). We collected 405 RNA-seq samples (Table S2), capturing 25 specific organs at different developmental stages, categorised into four major organs - leaves, roots, stems, and reproductive organs - for simplified comparison (Figure 1a)(Table S1). Our transcriptome assembly pipeline combined TRINITY and k-mer SOAPdenovo-Trans assemblies concatenated with EvidentialGene (Figure S2)^36,37^. We removed any potential non-fern mRNA contaminants and any sequences with aberrant GC content due to assembly artefacts, low transcripts per million (TPM) values, or sequence similarity higher to non-fern species than to ferns (see methods, Figure S2). The assembly yielded 30,000–100,000 coding sequences (CDSs) per species with high Benchmarking Universal Single-Copy Orthologs (BUSCO) scores (Figure 1b, Table S3) that rivalled the scores of the four sequenced genomes (Figure 1b, black arrows).

**Figure 1.**
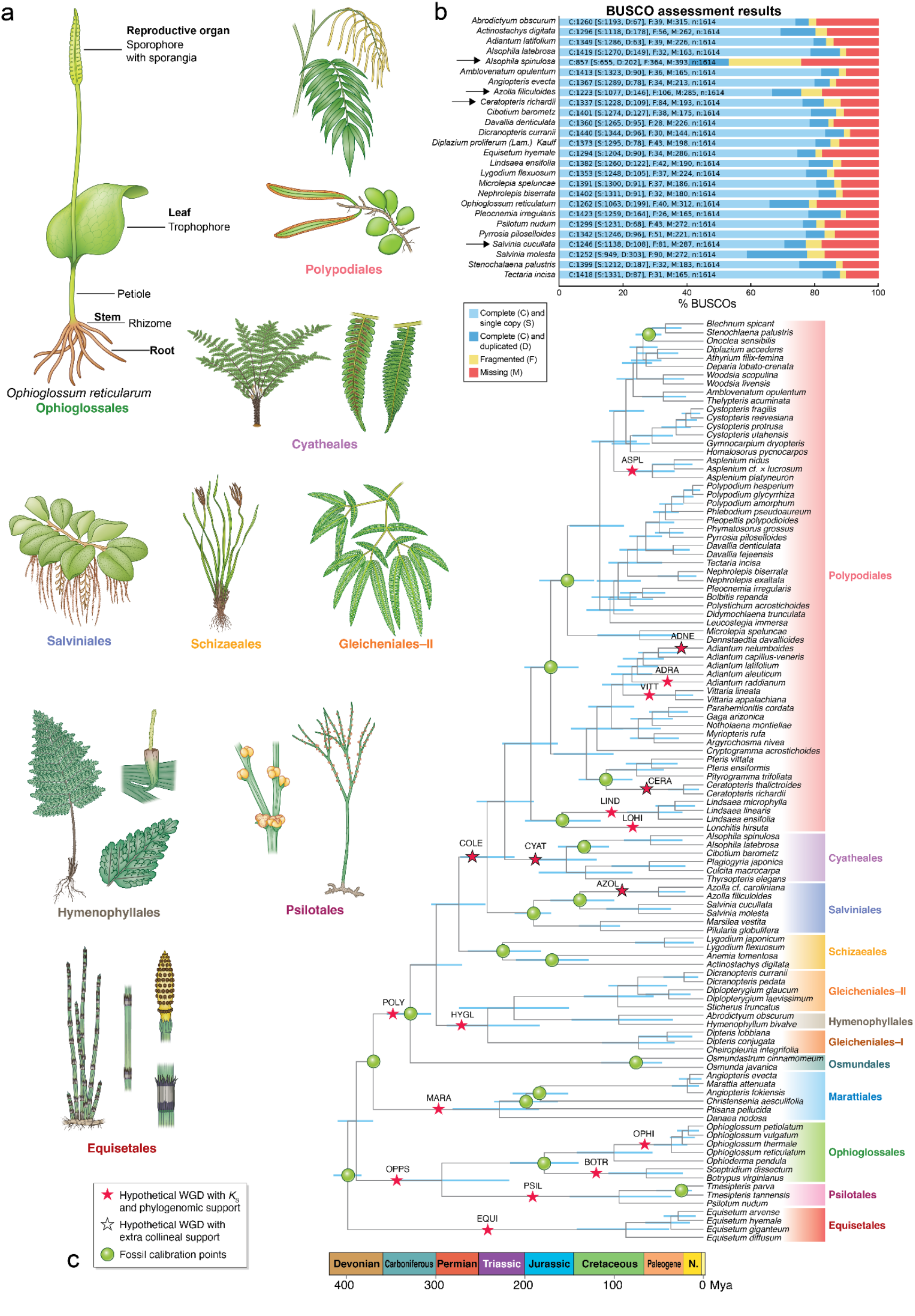
Sampling, transcriptome assembly, and species tree of major representatives of ferns. a) *Ophioglossum reticulatum* with sampled organs labelled, together with samples representatives of the other fern ordersb) Completeness of transcriptome assembly measured by Benchmarking Universal Single-Copy Orthologs (BUSCO). *Alsophila spinulosa, Ceratopteris richardii, Azolla filiculoides* and *Salvinia cucullata* have available genomes, while the remaining values are for the transcriptome assemblies reported here. C) The evolutionary timescale of Monilophyta based on the inferred consensus fern cladogram. The species tree shows the inferred consensus phylogenetic topology with branch lengths representing absolute divergence time estimated by Bayesian molecular dating analysis. The horizontal coordinates of each internal node denotes the posterior mean divergence time while the bars represent the 95% Highest Posterior Density. Hypothetical WGDs are indicated at corresponding phylogenetic nodes as red stars with four-letter identifiers. Black outline around a red star indicates events with additional collinear support. Fossil calibrations are indicated at corresponding phylogenetic nodes with green circles. The clade strips indicating affiliations at order level are shown as vertical bars with distinct colours. The geological timeline refers to the International Commission on Stratigraphy (ICS) v2023/09.

### Reconstruction of the evolutionary timeline of ferns

Given the standing phylogenetic discordance, we reconstructed a phylogenetic tree of 108 fern species (22 from this study, 7 sequenced genomes, and 79 from the 1000 Plant Transcriptomes Initiative (1KP) and other studies)^13,20,32–35,38–40^, covering the whole backbone of Monilophyta (Table S4, Supplemental Methods 1). Four datasets, each with a different outgroup (horsetails, seed plants, lycopods or bryophytes), were first used in nucleotide with three different methods including ASTRAL-Pro2 ^41^, concatenation-based method and STAG ^42^(Supplemental Methods 1) on the 107 ferns dataset and then reanalyzed in both nucleotide and peptide on the 108 ferns (adding the latest *Marsilea vestita* genome ^39^) using the favored method ASTRAL-Pro2. We recovered a well-supported fern backbone phylogeny in which Marattiales was inferred to be sister to Polypodiidae (i.e. leptosporangiate ferns) and Gleicheniales was inferred as a paraphyletic clade (Supplemental Methods 1, Figure SM5-6). A closer phylogenetic relationship between Marattiales and Polypodiidae than between Marattiales and Ophioglossidae was supported in all datasets with Local Posterior Probability (LPP) as 1.00, except the one with bryophytes as outgroup based on peptide alignment. Phylogenies derived from this dataset favored Marattiales and Ophioglossidae as sister groups, with LPP as 0.82 (Supplemental Methods 1, Figure SM6). The monophyly of Gleicheniales was not supported, whereas the Gleicheniales−II (Gleicheniaceae) was closer to Hymenophyllales than Gleicheniales−I (Dipteridaceae) in all datasets, except the one with bryophytes as outgroup based on nucleotide alignment, whose conflicting branching pattern was only supported by LPP as 0.46 (Supplemental Methods 1, Figure SM5).

In addition, these analyses provided strong support for a scenario wherein horsetails are a sister group to the last common ancestor of all the remaining ferns in the phylogeny inferred from every combinational setting of non-horsetails outgroups and methods (Supplemental Methods 1). We selected phylogeny derived from the nucleotide dataset with horsetails as outgroup using ASTRAL-Pro2 as the consensus tree and used in all our subsequent analyses (Figure 1c). This selection was based on the general consistency across datasets with varied outgroups, and on the ASTRAL-Pro2 derived phylogeny with STAG method (Supplemental Methods 1).

To estimate the absolute divergence time of the 108 ferns, we used Bayesian molecular dating under the independent rate and LG general amino acid substitution model ^43^ with 18 soft fossil constraints (indicated in Figure 1c as green dots, Table S5). The 95% Highest Posterior Density (HPD) and posterior mean of the stem ages of major fern clades was summarised (Supplemental Methods 1, Table S6). The resulting high-confidence fossil-calibrated tree thus resolved a long-standing discussion on fern phylogenetics.

### Identification of 18 separate whole genome duplication events in ferns

It has been proposed that ferns have a large number of chromosomes due to repeated rounds of whole-genome duplication (WGD)^44^. Here, we used *K*S-age distributions and phylogenomic methods to unveil remnants of ancient WGDs ^45^. In total, we found support for 18 hypothetical WGDs within the backbone of ferns, of which five with attained collinear support (red stars, Figure 1c, see Supplemental Methods 1). Seven of the WGDs were found within Polypodiales, two in Ophioglossales, and one in each of the lineages of Equisetales, Psilotales, Marattiales, Salviniales and Cyatheales, together with four shared by more than one order. WGDs were previously identified in other studies with different phylogenetic locations ^28,46^. We reevaluated possible scenarios thereof and proposed WGDs with both *K*S and phylogenomic support after correcting rate variation and taking into account the uncertainty in gene tree and gene tree-species tree reconciliation (Supplemental Methods 1). Next, we tested the species richness and genome size as a function of the number of ancient WGD events shared by different major clades (excluding Polypodiales due to numerous nested WGD events therein), and observed a significant positive correlation between the number of ancient WGD events and the number of species in a lineage (Figure S3, adjusted R^2^ = 0.41, p-value = 0.02). Conversely, we did not observe any significant correlation between the number of ancient WGD events and genome size (Figure S3).

### Diversity and conservation of fern gene functions and expression patterns

To investigate predicted gene functions within ferns, we conducted a phylostratigraphic analysis encompassing one glaucophyte, seven chlorophytes, three bryophytes, two lycophytes, 26 ferns (22 fern transcriptomes and four genomes), two gymnosperms, and six angiosperms (see methods). The 47 species are used to study the gain and losses of gene families across Archaeplastida, by assigning the orthogroups to nodes ranging from node 1 (the earliest ancestor of Archaeplastida) to node 13 (the ancestor of Polypodiales)(Figure 2a). The nodes are based on the fern phylogeny tree shown in Figure 1c and known relationships between Archaeplastida ^47^.

**Figure 2.**
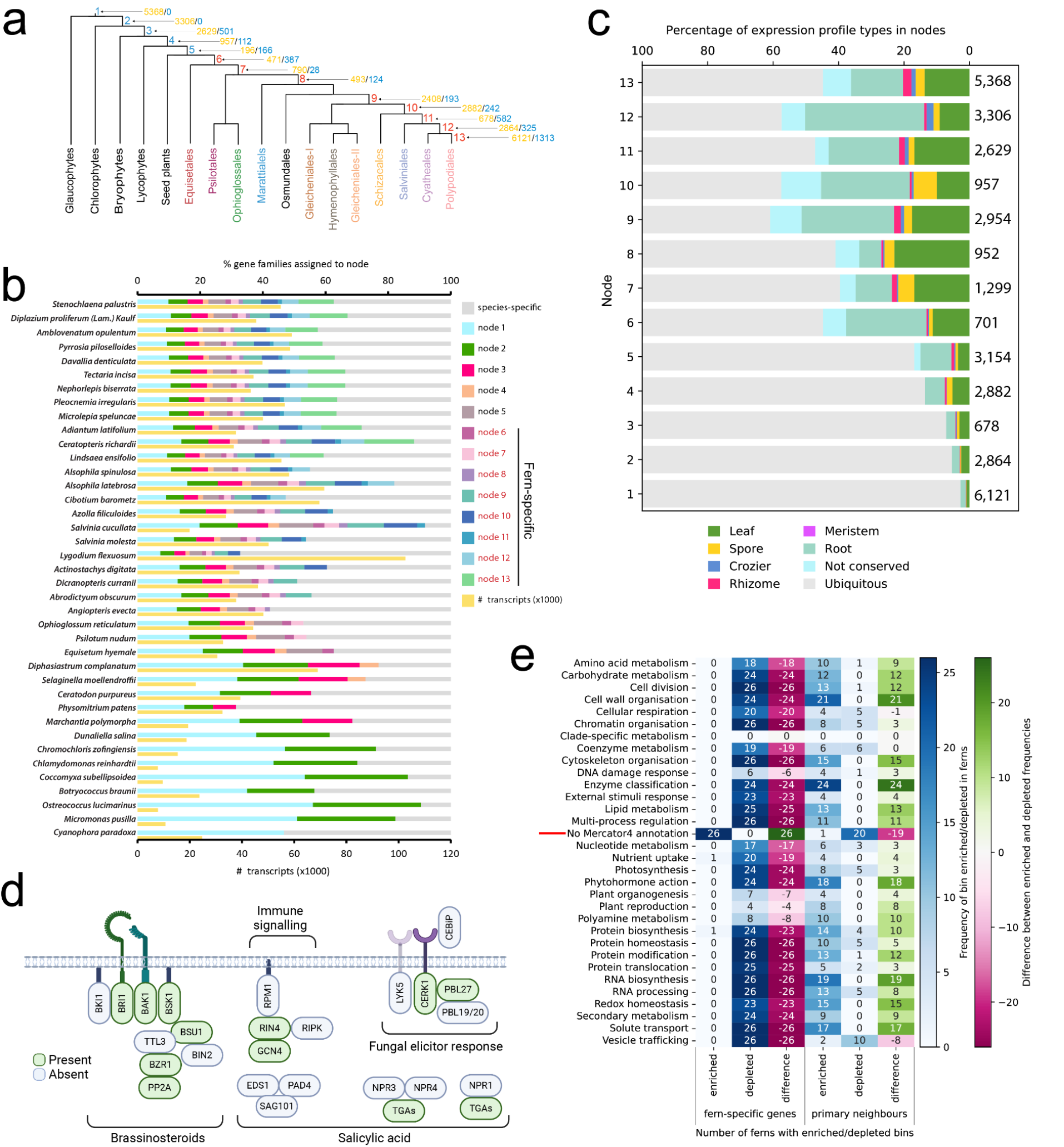
Gene functions in ferns. a) Division tree of Archaeplastida. Leaves represent orders, while node numbers correspond to the phylostrata. The orange and blue numbers indicate the number of gains and losses of orthogroups, respectively. b) Stacked bar plot showing the percentage of gene families belonging to a phylostrata (node). Species- specific gene families comprised genes from only one species and were not assigned to nodes. c) Percentage of gene families identified as organ-specific for each node. Numbers on the right side of the plot indicate the number of orthogroups per node. d) Examples of signalling components present (green shapes) or absent (grey shapes) in ferns. e) Clustered heatmap showing enrichment and depletion of biological processes of fern-specific genes and primary co-expression neighbours in 26 (22 from this study and four sequenced genomes) fern species analysed. The left column indicates the biological processes defined by Mapman bins. The scores in the ‘enriched’ and ‘depleted’ column indicate in how many of the 26 species fern-specific genes are significantly (BH adj.p < 0.05), connected or disconnected to genes belonging to a specific bin, respectively. A higher value in difference (# enriched - # depleted) indicates overall enrichment, while lower values indicate overall depletion for respective bins in the 26 ferns.

Orthogroup gains, a measure of new gene family acquisition, were highest in the early stages of algae evolution (nodes 1 and 2 gained 5368 and 3306 orthogroups, respectively) and when plants colonized the land (node 3, 2629 gained orthogroups)(Figure 2a). However, a substantial number of orthogroup gains and losses were also observed within the different fern lineages (e.g., 2408 gained and 193 lost orthogroups in node 9, Figure 2a).

The analysis further revealed that ∼50% of fern gene families are fern-specific (Figure 2b nodes 6–13 in red, Table S7). For example, >50% of gene families in *Stenochlaena palustris* belong to nodes 6–13 (fern-specific nodes 6–13 in red, Figure 2a, Table S7), suggesting that the fern lineage has evolved genes with novel, unexplored functions. We analysed sequences of 25 orthogroups comprising at least ten fern species. Representatives of 17 orthogroups show no significant sequence or structural similarity to non-fern species (Figure S4), with eight of them being disordered (few to no secondary structures). These results show that >50% of gene families in ferns represent novel, uncharacterized proteins, indicating that studying ferns will likely provide new insights into plant biology and evolution.

To understand how gene age correlates with gene expression specificity, we identified organ-specific genes with specificity measure (SPM) analysis (distributions of SPM values are shown in Figure S5, expression profiles of organ-specific genes are shown in Figure S6, Table S8)^47^. Genes belonging to older nodes (nodes 1–4) were less organ-specific (<20%) than younger nodes (nodes 5–13, 30–60%, Figure 2c). Younger genes (nodes 5–13) had specialised functions in a particular organ, with roots having the highest number of specifically expressed genes (Figure 2c). Furthermore, species-specific genes also tended to show a less ubiquitous, more organ-specific expression (Figure S7), indicating an overall negative association between gene age and organ specificity, which is in line with similar observation in land plants ^47^.

Finally, we investigated whether organ-specific genes are conserved across ferns and other land plants. While many organs express significantly similar sets of genes across ferns and other land plants, we observed a clear difference between fern and seed plant transcriptomes (Figure S8). Not surprisingly, the organ-specific gene sets of seed-containing plants show higher mutual similarity to those of other seed plants (Figure S8, blue box), while those of ferns show the highest similarity to those of other ferns (Figure S8, green box).

### Ferns lack several genes essential for hormonal signalling, defence and development in angiosperms, indicating their unique developmental and environmental strategies

Terrestrialization and the evolution of seeds and flowers required the evolution of many biological functions ^48^, which is readily visible when comparing gene inventories of algae, land, seed and flowering plants (Figure S9, gene function completness indicated by darker cells). We used MapMan sequence-based annotations and compared the gene function repertoire of ferns and model angiosperms and observed the absence of several components in ferns (missing functional categories indicated with red text in Table S9, Supplemental Methods 2).

#### Hormone signalling

Missing components include abscisic acid regulation, perception, and transport, auxin methylation-based degradation, brassinosteroid signalling (Figure 2d)^49^ and degradation, cytokinin degradation and transport, and degradation of gibberellins and jasmonic acid and their transport genes.

For example, several components of the salicylic acid (*SAG101, EDS1, PAD4*, Figure 2d)^50^ and strigolactone signalling pathways were missing (Table S9), as is the degradation component of the former hormone. To test this further, we investigated the presence of canonical NPR domains (NPR1-like C superfamily, BTB/POZ NPR plant domain or BTB/POZ superfamily, and an ANKYRIN domain) in our fern transcriptomes. Of the 26 ferns we studied, 22 had at least one canonical NPR (Supplemental Methods 2, Figure SM7). This is consistent with previous evidence that the duplication of NPR1/3/4 happened sometime during angiosperm diversification, long after the split between flowering plants and ferns ^51^. The SAG101/PAD4/EDS1 module, on the other hand, appears to be a more recent invention of flowering plants, as it is mainly absent in non-seed plants (Supplemental Methods 2, Figure SM8).

Further, we investigated perception and downstream signalling of jasmonic acid (JA), focusing on the JA receptor COI1 and the JAZ transcriptional repressors. Both COI1 and JAZ candidates are encoded in fern transcriptomes. While *Arabidopsis thaliana* and *Marchantia polymorpha* both have only one copy of COI1, ferns show several gene duplication events, some of which are species-specific, and some of which appear more ancient (Supplemental Methods 2, Figure SM9). The current evolutionary model of JA perception is that the COI1 ligand switched from *dn*-cis-OPDA to JA-Ile in the ancestor of vascular plants ^52^. The radiation of COI1 in ferns, however, suggests functional divergence in jasmonate perception, possibly complicating its evolutionary history. This highlights ferns as a key lineage for further functional investigation to understand the evolution of plant immunity.

#### Secondary metabolism

Phytochemical studies on ferns have revealed that they contain a wide range of secondary metabolites, many of which are function herbivore defense and show bioactive properties ^9^. For secondary metabolite pathways associated with biotic interactions, we observed that multiple genes known to act in the flavonoid biosynthesis pathway were missing in all fern species analysed (Table S9), agreeing with previous datasets on the evolution of red pigmentation in land plants ^53,54^. Yet, ferns are able to synthesise flavonoids ^55^.

#### External stimuli response

Plants have evolved elaborate signalling and response pathways to cope with the changing environment. For several of these pathways, we observed that orthologs of phototropin-mediated receptors, all CO2 sensing and signalling components, and many gravity-sensing proteins were absent in our fern transcriptomes (Table S9). Genes known to be essential for sensing and responding to temperature in flowering plants were present in ferns, but acquired thermotolerance factors were missing. Other missing proteins include those involved in several pathogenesis-related processes, such as pattern- and effector-triggered immunity (16 out of 36 factors)(Figure 2d), WRKY33- dependent immunity, pathogen polygalacturonase inhibitors, and basic chitinases. While some components of symbiosis pathways are present in our fern transcriptomes (Table S9), many factors are absent, such as mycorrhizal response genes and transporters.

#### Transcript control and modification

Several components controlling mRNA and protein levels are also absent, such as more than half of the subgroups of MYBs and most REMs. Organellar RNA processing is lacking plastidial and mitochondrial CFM-type splicing factors, a majority of mitochondrial RBA splicing components (>20), C-to-U RNA editing (>50 factors), and mRNA stabilisation and deadenylation factors. For protein homeostasis, we found only class C-I and C-II small HSP holdase chaperones, while the ten other classes were absent (C-III to ER), together with E3 ubiquitin ligases from groups IV and V.

#### Reproduction and organ development

Not surprisingly, our transcriptomes indicate that ferns differ from flowering plants in their gene inventories related to reproduction. Ferns lack genes associated with anther dehiscence (*PCS1, NST1/2, MYB26*), pollen aperture formation (*INP1/2*), pollen tube growth (except *GEX3*), embryo axis formation (except *ATML1*), endosperm formation (exception *GLAUCE*) and seed formation and dormancy (Table S9). On the other hand, ferns contain nearly all male gametogenesis (e.g., *DUO1/3, DAZ, APD*) and exine (*ROCK/TEX2, DEX1, NEF1*) formation factors, stamen (*TPD1, EMS1, JAG*) and tapetum (*DYT1, TDF1*) regulators and most factors important for female gametophytes (*AMP1, CYP78a, RKD, MAA3*) but lack genes essential for central cell formation (Table S9). Surprisingly, while ferns are seedless, they contain most genes important for seed maturation and globulins.

Interestingly, most flower formation photoperiodic and autonomous promotion pathway genes are present in ferns and bryophytes. However, as expected, most genes important for floral transition are missing (*FRIGIDA, FRL1/2, FES1*), except *FRI-C* effector complex genes, floral meristem identity (*LMI2, AP1/3, PISTILLATA, SEPALLA*), and morphogenesis (*BLR, ETT*). This suggests that the flower formation pathways have other roles in ferns, possibly linked to photoperiodic response, developmental timing, sporulation control, or other process.

For organ development, ferns lack several key genes essential for leaf adaxial and abaxial polarity, guard cell formation, and stomatal density (Table S9). Their root developmental programs are likely also different from flowering plants, as they lack the entire MYB-bHLH-WD40 transcriptional regulatory module, and genes controlling columella apical meristem (*WOX5, FEZ, SMB, BRN1/2*), and endodermis meristem regulation and signalling (*SHR, SCR, KOIN, IRK*). More than half of Casparian strip factors are missing, and nearly all vascular system formation factors (only 2 out of 14 transcription factors present).

Taken together, the analyses shown in Figure 2a-e provide further support for the presence of unique growth, development, and survival strategies in ferns, and suggests that additional research on them is worthwhile.

### Co-expression-based prediction of gene functions in the fern lineage

The presence of >50% of fern-specific orthogroups (Figure 2b) indicated that ferns might have evolved as-yet unknown gene functions on a massive scale. To investigate whether the fern-specific genes can be annotated by sequence-similarity approaches, performed an enrichment analysis of their biological functions. We observed enrichment for MapMan bin containing uncharacterized genes (‘No Mercator4 annotation’, red line, Figure 2e), and a depletion of bins related to known biological processes. This indicates that sequence similarity approaches cannot infer the functions of most fern-specific genes.

Gene co-expression networks can reveal the functions of non-annotated genes based on the guilt-by-association principle ^56^, where genes with similar expression profiles tend to be involved in the same biological process. To predict the functions of fern-specific genes without relying on sequence similarity to genes with known functions, we calculated the functional enrichment of their direct neighbours in the co-expression networks. Interestingly, fern-specific genes are significantly co-expressed (> 17 fern species) with biological processes such as ‘Cell wall organisation’, ‘Enzyme classification’, ‘Phytohormone action’ and ‘RNA biosynthesis’ and moderately co-expressed (> 10 fern species) with ‘Cell division’, ‘Cytoskeleton organisation’, ‘Lipid metabolism’, ‘Multi-process regulation’, ‘Protein biosynthesis’, ‘Protein modification’, and others (Figure S10, Figure 2e). This indicates that these genes are involved in most biological processes in ferns, especially cell wall, development and new metabolic pathways.

### S lignin has evolved independently in the fern lineage

Lignin is a complex phenolic polymer that forms essential structural materials in the support tissues of vascular plants. Importantly, this polymer also confers hydrophobicity to xylem vessels, allowing water transport from roots to leaves and enabling plants to grow out on land. Lignin is primarily composed of three monolignols: *p-*coumaryl, coniferyl and sinapyl alcohols, which are named *p-*hydroxyphenyl (H), guaiacyl (G) and syringyl (S) units when incorporated into the polymer ^57^. Several studies, including one focused on ferns ^58^, suggest a complex evolutionary history that may include independent evolutionary paths for lignin synthesis, particularly the S units, among different plant lineages ^59^. However, testing this was difficult without additional genomic information. Our fern transcriptome datasets provided an opportunity to explore the evolution of lignin across the entire fern family.

We first characterised the presence and sites of deposition of the different lignin units in nine ferns from three orders: Equisetales, Cyatheales and Polypodiales, using a simple staining procedure. We stained cross-sections of stems and petioles with Phloroglucinol-HCl, which reacts with coniferaldehyde residues of lignin to generate a red condensation product ^60,61^. All nine ferns showed the presence of lignin. However staining was not or poorly discernable in vessels of *Equisetum hyemale* and *Adiantum latifolium* (Figure 3a, Figure S11). In many species, lignin was mostly found in the subcortical sclerenchyma or outer layers of the metaxylem tissues, as expected (Figure S11). Interestingly, Mäule staining on stem cross-sections indicated the presence of S-units in *Pleocnemia irregularis* and *Stenochlaena palustris* ^62^ (Figure 3a, Figure S11), indicating that this subunit predominantly found in angiosperms ^63^, is also widespread in ferns.

**Figure 3.**
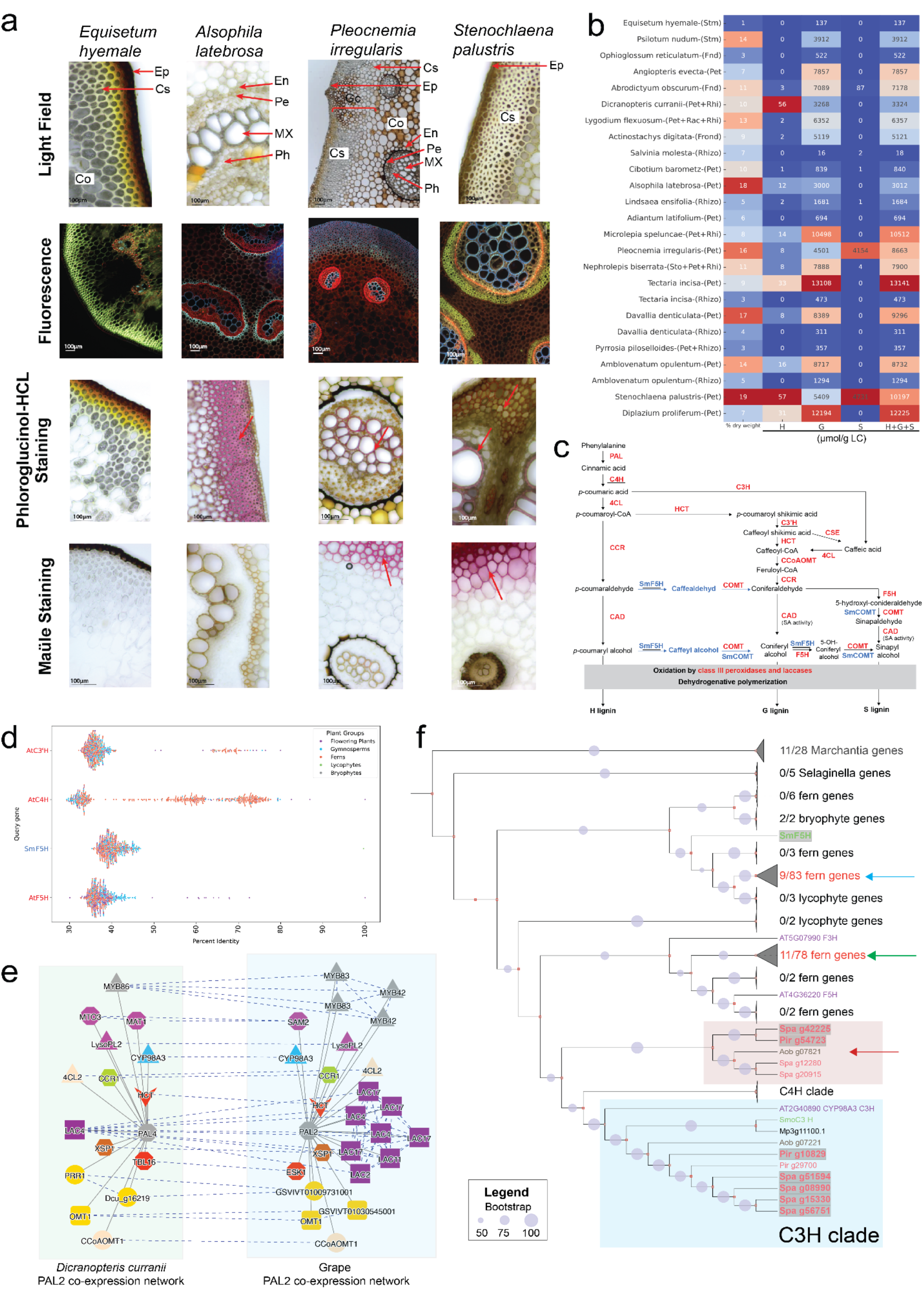
Lignin analysis of ferns. a) Sections of stems (Equisetum) and petioles (Alsophila, Pleocnemia, Stenochlaena). Red arrows with labels indicate cortex (Co), cortical sclerenchyma (Cs), endodermis (En), epidermis (Ep), metaxylem (MX), pericycle (Pe) and phloem (Ph). Red arrows without labels indicate stained cell walls. b) Percentage of lignin (1st column), H, G, S (2nd-4th) and H+G+S (5th) thioacidolysis lignin units. c) The H, G, and S lignin unit biosynthesis pathway for angiosperms (red text) and lycophytes (blue text). Intermediate metabolites are indicated by black text, while the grey box contains enzymes involved in lignin polymerization. d) BLAST scores (x-axis) of AtC3H, AtC4H, SmoF5H and AtF5H against the translated transcriptomes found in the CoNekT database. Each point represents a protein. e) Comparative co-expression network analysis of *PAL* genes from grape and fern *Dicranopteris curranii.* Nodes represent genes, solid edges connect co-expressed genes, while dashed edges connect orthologs. Coloured shapes represent different orthogroups, while the gene names are based on the best BLAST hits to *Arabidopsis thaliana*. f) Phylogenetic tree of land CYP450s of *P. irregularis* (genes starting with Pir), *S. palustris* (Spa), *A. obscurum* (Aob), *Selaginella moellendorffii* (Smo), *Marchantia polymorpha* (Mp). The Arabidopsis (AT) lignin-related C4H, C3’H and F5H and flavonoid-related F3H are included.

Given these results, we further determined lignin content and structure within all 22 fern species, using the CASA method ^64^ and thioacidolysis followed by Gas Chromatography-Mass Spectrometry (GC-MS) ^65,66^, respectively (Figure 3b). Overall, we observed a large variability in total lignin content and the different units among the ferns. Not surprisingly, *Equisetum hyemale* showed the lowest CASA lignin content (1 %), and *Stenochlaena palustris* the highest (19 %)(Figure 3b). Most ferns contain lignin composed of G units (130 - 13000 µmol/g of CASA lignin), while H units are less abundant and, in some cases, not detectable (0 - 57 µmol/g). Interestingly, we observed substantial differences in the lignin content of multiple organs when analyzed. For example, H units were detectable in petioles but not in rhizomes of *D. denticulata* and *A. opulentum* (Figure 3b, Table S11). We observed S units in high quantities in *P. irregularis* and *S. palustris* (>4000 µmol/g), medium quantities in *A. obscurum* (86.9 µmol/g CASA) and minute but detectable quantities in *S. molesta, Cibotium barometz, Lindsaea ensifolia, Nephrolepis biserrata* (<5.0 µmol/gCASA).

Next, we set out to identify the biosynthetic pathways of lignin, focusing on S units. To analyze the pathways and make the fern gene expression data easily accessible, we uploaded the expression data for the 22 ferns to our CoNeKT database (https://conekt.sbs.ntu.edu.sg/)^67^, upgrading the database to comprise 39 species, including angiosperms, lycophytes, bryophytes and algae. S unit synthesis evolved independently in the lycophyte *Selaginella* and angiosperms ^68^, illustrated in Figure 3c. Unlike angiosperms, which require *p-*coumarate 3-hydroxylase (C3’H) and ferulate 5- hydroxylase (F5H) to make S units (Figure 3c, black arrows), *Selaginella* utilises a multifunctional F5H that skips several steps of the pathway to make caffealdehyde and caffeyl alcohol, which can then be utilised to make G and S lignin (Figure 3c, blue text)^69^. Blasting *AtC3H* and *AtC4H* (Cinnamate 4-hydroxylase) against all species proteomes in the CoNekT database (https://conekt.sbs.ntu.edu.sg/blast/), showed identity scores of >60% for ferns (Figure 3d, orange points), indicating that ferns likely contain C3’H and C4H enzymes (Table S12). However, *AtF5H* and *SmF5H* showed only low sequence identity to fern proteomes (∼40%), which according previous studies indicates an absence of known F5H enzymes in ferns ^70^.

To identify candidate fern F5H enzymes, we took advantage of the observation that lignin biosynthetic genes tend to be tightly coexpressed and that these relationships are conserved even across large evolutionary distances ^71^. Indeed, comparing the co-expression networks of *PAL* genes from fern *Dicranopteris* and angiosperm *Vitis vinifera* (grape, Vitaceae) revealed many of the expected enzymes and a CYP98A3-like gene that could likely represent *C3H* (Figure 3e, query gene *Dcu_g01768*, co-expression networks of the lignin genes are in Supplemental Data 1). Furthermore, most of the fern lignin biosynthetic genes are co-expressed with at least two other relevant enzymes (Figure S12a), and the co-expression networks can suggest the unknown components (Figure 3f), including transcription factors and CYP450 enzymes (Figure 3f, Figure S12). To suggest the identity of fern F5H enzymes, we first performed phylogenetic analysis of all CYP450s of S lignin-producing ferns Pir, Spa, Aob, angiosperm *Arabidopsis*, lycophyte *Selaginella* and included the outgroup bryophyte *Marchantia* that does not produce S units (Figure 3g). We then indicated which CYPs are co-expressed with at least one lignin enzyme. As expected, *C3H* genes are co-expressed with the other lignin biosynthetic enzymes (co-expressed genes indicated with grey boxes, Figure 3g). However, we observed several clades in the tree that likely emerged independently in ferns and contained groups of co-expressed CYP450 enzymes (Figure 3g, indicated by red, green, and blue arrows). These enzymes comprise prime candidates for the discovery of F5H enzymes in ferns.

### Members of the Polypodiales contain a non-canonical cell wall sugar

To further understand the evolution of fern cell walls, we carried out a monosaccharide composition analysis using Gas Chromatography on the 22 ferns that were part of our transcriptomic study (Table S13). The most abundant sugars were glucose (a building block of cellulose, mixed-linkage glucans, xyloglucans), mannose (mannans) and xylose (xylans)(Figure 4a). Less abundant sugars were rhamnose (pectic rhamnogalacturonan I, arabinogalactan-protein), fucose (rhamnogalacturonan II, xyloglucan, arabinogalactan-protein), arabinose (hemicellulose arabinoxylan, rhamnogalacturonan I and II, arabinogalactan-protein) and galactose (rhamnogalacturonan I, hemicellulose galactomannans, arabinogalactan-protein). The proportions of various sugars changed among species and among different organs of the same species, which is in line with previous observations ^72^. For example, the *T. incisa* rhizome exhibited a higher proportion of glucose than the petiole of the same species, and higher than rhizomes in of *Davaillia denticulata* and *Ambloventanum opulentum* (Figure 4a). In addition to these sugars, we also observed trace amounts of methylated rhamnose (3*O*-MeRha*p*), a known sugar found in arabinogalactan proteins in ferns ^73^, in all species except for *Psilotum nudum* and *Dicranopteris curranii* (Figure S13),

Interestingly, we detected an unknown peak from samples derived from three species, *T. incisa*, *A. opulentum* and *D. proliferum* (Figure 4a, red bars). Because this peak was not observed with our common standards during Gas Chromatography analysis (data not shown), it likely represented a novel sugar. Since initial GC-MS analyses suggested the sugar to be a methylated hexose (data not shown), we synthesised a panel of methylated sugars (Figure S14). Out of the six methylated sugars, only 2-*O*-Methyl-D-glucopyranose (2O-Me-Glc*p*) showed identical retention time and mass spectrum to the unknown sugar (Figure 4b, Figure S15), indicating that these three species of ferns produce a novel sugar. While methylated sugars such as 4-*O*-D-Methyl-Glucuronic Acid are present in plant cell walls ^74^, this is to our knowledge the first report of 2O-Me-Glc*p* and warrants further study.

**Figure 4.**
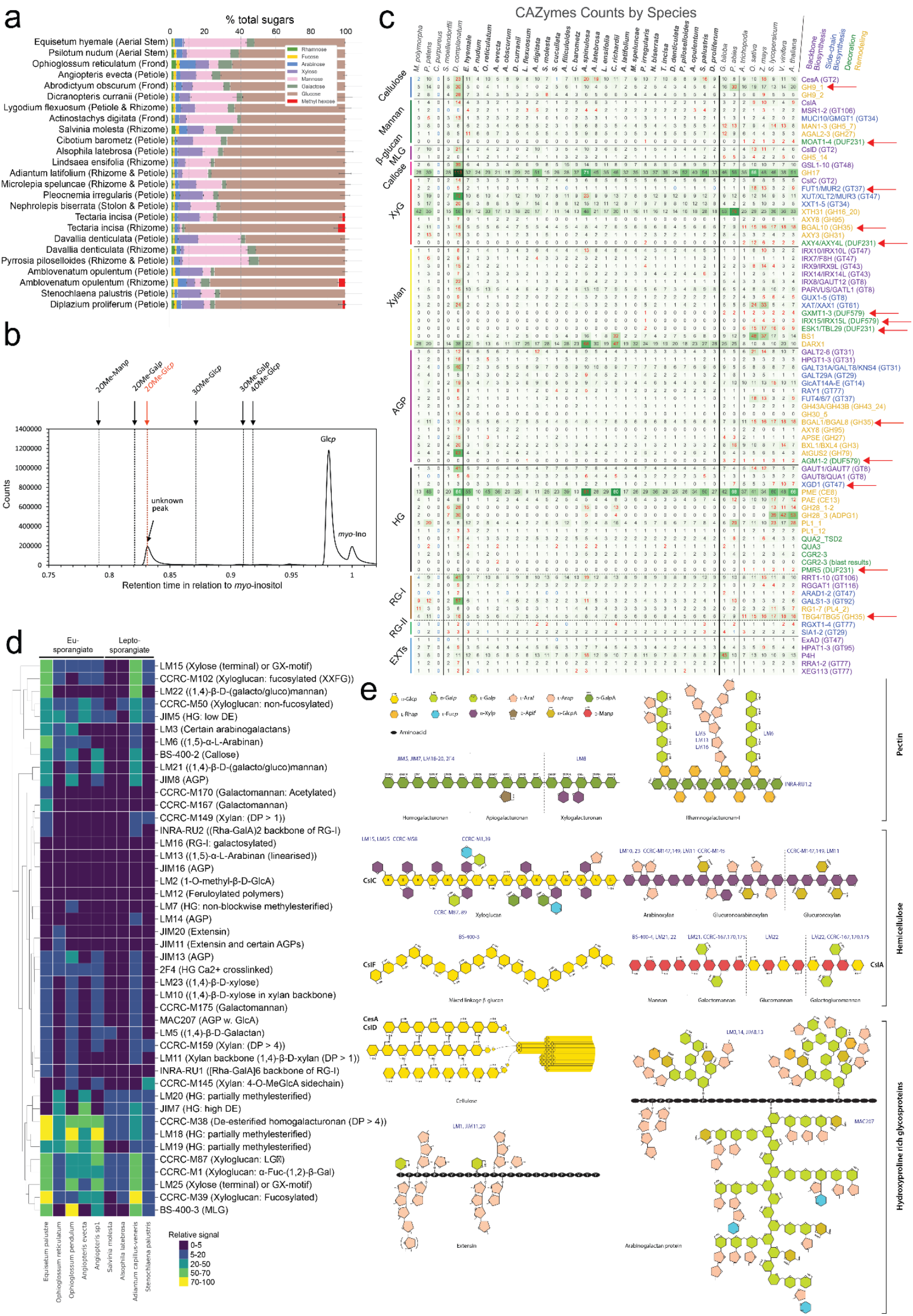
Polysaccharide analysis. a) Percentage of total neutral sugars estimated by GC-MS. The sugars are rhamnose, fucose, arabinose, xylose, mannose, galactose and glucose. b) GC spectra of the unknown peak from c) The number of orthogroups involved in cell wall biosynthesis in land plants. Columns represent species, while rows correspond to a given gene family. The rows are further divided into different polysaccharide classes, separated by horizontal lines. Red and blue numbers indicate that a given species contains significantly more/less genes than others (adjusted p-value < 0.05). Darker colors of boxes indicate more gene copies in each row. The red arrows indicate rows which are particularly depleted in ferns. Ferns are indicated with bold names and thick black lines. d) Relative epitope abundance for five fern species quantified by Comprehensive Microarray Polymer Profiling (CoMMP). Rows indicate the different species and organs, while columns represent the obtained signal from the different antibodies. The colours of the cells correspond to the signal strength. e) Schematic drawing of the major cell wall polysaccharides. Each colour-coded shape represents a sugar or amino acid. The antibodies binding to a respective epitope are coloured with blue letters, while the black bold letters indicate the biosynthetic enzymes.

### Ferns contain most but not all cell wall polysaccharides of angiosperms

We next performed a large-scale comparative analysis of Carbohydrate-Active enZYmes (CAZymes) in land plants (Figure 4c)^75^. The CAZyme database contains genes involved in cell wall biosynthesis, allowing us to compare similarities and differences of ferns and other land plants. To do this, we calculated with species contain significantly (adjusted p-value <0.05) more (red numbers) or less (blue numbers) than the other species. Ferns contain fewer xyloglucan-related gene families involved in remodelling (*BGAL10*)^76^, fucosylation (*FUT1*) and no genes involved in O-acetylation (*AXY4*)^77^ (Figure 4c, red arrows)^78^. Although the number of FUT1 homologs are fewer in ferns, the evolutionary history of GT37 sequences was shown to be more complex when it comes to substrate specificity ^79^. For xylans, ferns showed a near absence of genes involved in methylation of glucuronic acid in glucuronoxylan (*IRX15*, *GXMT1-3* ^74^) and xylan acetylation (*ESK1*)^80^. Homogalacturonan pectins showed fewer fern genes involved in xylogalacturonan synthesis (*XGD1*)^81^ and galacturonan acetylation (*PMR5*)^82^. Finally, rhamnogalacturonan I pectins showed fewer fern genes involved in remodelling (*TBG4*)^83^. A similar analysis of hydroxyproline-rich glycoproteins did not reveal significant absences of these proteins in ferns (Figure S16). Overall, ferns tend to contain a lower number of genes per family and do not have the *DUF579* (sugar methylesterification)^84^ and DUF231 (sugar acetylation)^85^ gene families (Figure 4c).

To directly compare the polysaccharide inventories of angiosperms and ferns, we performed a Comprehensive Microarray Polymer Profiling (CoMPP)^86^, where we probed 102 cell wall extracts from nine ferns from six fern orders, including 36 organs at different developmental stages, with 48 antibodies targeting different cell wall epitopes The epitopes recognised by the antibodies are given in Table S14.

Our analysis revealed the relative abundance of cell wall polysaccharides (Table S15), which showed that different organs from the same species tend to have similar cell wall composition (Figure S17a) and the polysaccharide profiles within species tend to be more correlated than across species (Figure S17b).

We found that Eusporangiate ferns were generally richer in easily extracted polymers than leptosporangiate ferns, with Adiantum as a notable outlier (Figure 4d). Surprisingly, while mixed-linkage *β*-glucan (MLG) was only reported so far in Equisetum ^87,88^, we observed a clear MLG signal outside of Equisetidae in *O. pendulum, Angiopteris sp1, A. capillus-veneris* and *S. palustris* (Figure 4d). For pectins, we observed a high abundance of homogalacturonan at different grades of methyl esterification (antibodies CCRCM38, LM18, LM19, LM20, JIM5, JIM7), but low signal from RG-I backbone (INRA- RU1, INRA-RU2), galactosylated RG-I (LM16) and arabinan (LM13). For hemicelluloses, we observed a strong signal for xyloglucan (CCRC-M87), both fucosylated (CCRC-M102, CCRC-M1, CCRC-M39) and non-fucosylated (CCRC-M50), suggesting that xyloglucan might be a quantitatively important hemicellulose, which contrasts with a previous study reporting mainly mannan-rich cell walls ^89^. We also observed signals from xylan (CCRC- M154, LM11, CCRC-M159, LM10, LM23) and galactomannan (CCRC-M175) and various mannan-containing polysaccharides (LM22, LM21), with galactomannan showing signal only in *E. palustre* (CCRC-M170, -M167). The weakest signals were observed for epitopes in arabinogalactan-proteins (AGPs) and extensins, as only a few antibodies gave moderate signals (JIM8, MAC207, JIM13). Other antibodies showed weak signals (AGPs: JIM16, LM2, LM14, extensins: JIM20, JIM11). Finally, feruloylated polysaccharides that crosslink with arabinan and galactan residues of cell wall pectin via ester bonds ^90^ showed no signal (Figure 4d). Taken together, these results indicate that fern and angiosperms share most of the polysaccharides and their biosynthetic enzymes, but ferns might lack certain sugar modifications and AGP structures found in flowering plants.

### Evolution of the cellulose synthase superfamily in Archaeplastida

In addition to lignin, cellulose is one of the major load-bearing polymers. Angiosperms contain primary and secondary cell walls enriched in cellulose, which are biosynthesised by CESA1,3,6 and CESA4,7,8 in *Arabidopsis thaliana* ^91^. The CESA complexes are arranged in hexameric complexes called rosettes in angiosperms ^92,93^, or as linear terminal complexes in bacteria ^94,95^. *Selaginella* is the latest diverging plant known to possess both CesA hexameric complexes and CesAs of the type that forms linear complexes in bacteria ^96^. The plant kingdom has also evolved cellulose synthase-like (*CSL*) genes to produce other polysaccharides ^97^, such as mannans (*CSLA*)^98^, glucan chain of xyloglucan (*CSLC*)^99^, cellulose in tip growing cells (*CSLD*)^100,101^ and mixed- linkage glucans (*CSLF*)^102^. However, the evolution of the CESA superfamily is not well understood in ferns.

A phylogenetic analysis of the *CESA* superfamily built from algal and land plant protein sequences showed that ferns contain both linear (*CESA* linear) and hexameric (*CESA1/3/10* and *CESA6*) *CESA* genes (Figure 5a, Figure S18). The *CSLA*, *CSLC* and *CSLD* families were found in all land plants, including ferns (Figure 5a). The *CSLB* and *CSLG* families are only found in seed plants ^103^, but ferns contain one clade of genes that is likely ancestral to the two families (Figure 5a, green arrow). We also observed that the angiosperm secondary cell wall enzymes *CESA4,7,8* and the fern *CESAs* do not form a monophyletic group (Figure 5a, black arrows), indicating that ferns either lack CESA4, 7, 8 or have evolved versions that no longer form clear clades with them. Conversely, ferns form two distinctive groups with Arabidopsis *CESA6* and *CESA1,3,10* (Figure 5a), suggesting that the ancestor of ferns and seed plants contained two *CESAs* that gave rise to *CESA6-like* and *CESA1-like* clades. The phylogenetic tree revealed four independent duplication events of the *CESAs* within ferns (Figure 5a, light blue arrows), suggesting that ferns have likely evolved cell walls with properties distinctive from flowering plants.

**Figure 5.**
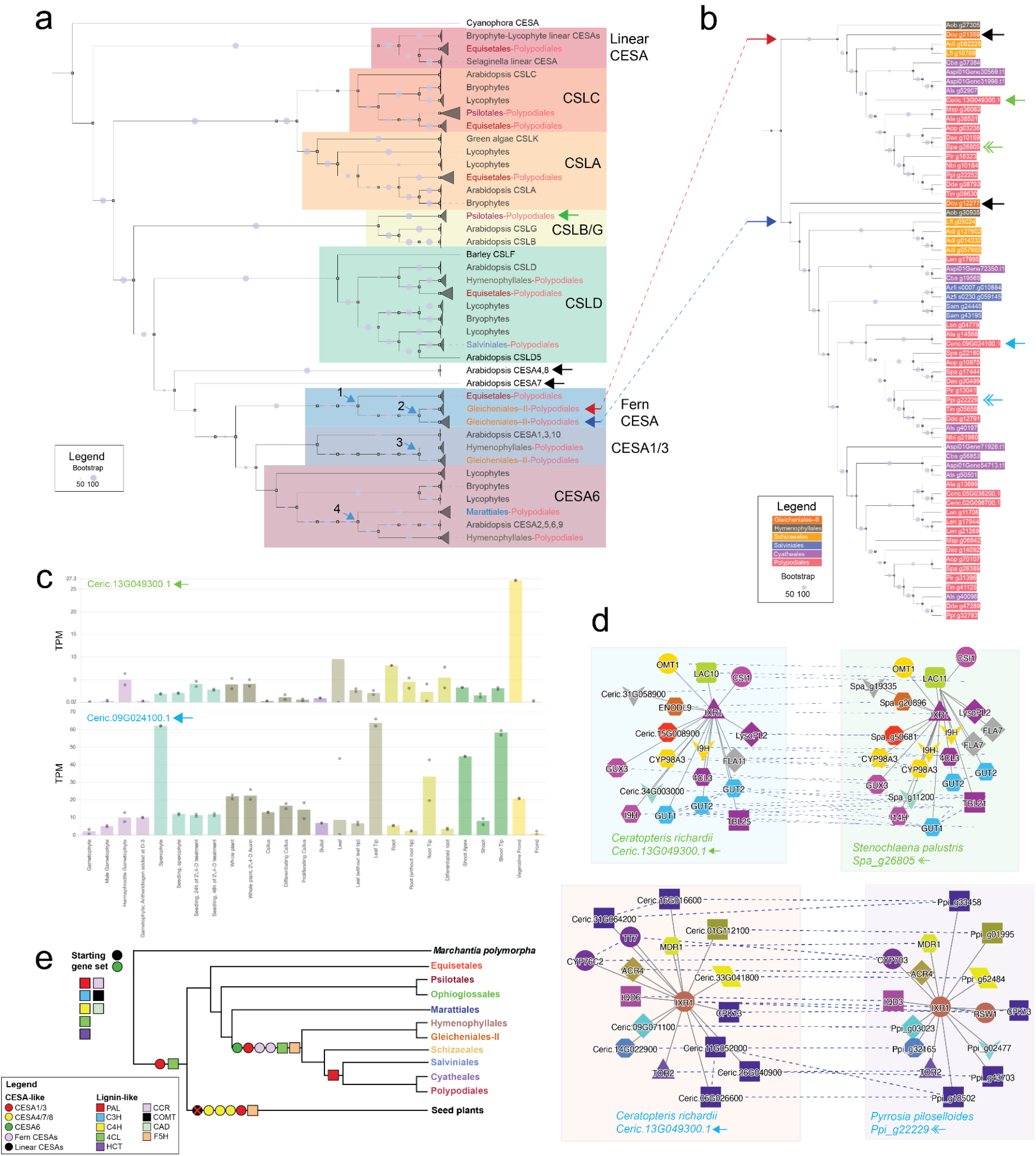
Independent duplication of cell wall-related modules in ferns. a) Maximum likelihood cellulose synthase superfamily gene tree. Gray triangles indicate collapsed clades. The arrows indicate the discussed clades and duplication events. The grey circles indicate the bootstrap support of the nodes, while the colored text represents the different fern clades. b) Closeup on the fern-specific clades. The fern orders are colour-coded, while the blue and green arrows indicate the discussed genes. c) CoNekT expression profiles of two CESA genes from *Ceratopteris richardii*, where the x-axis indicates organs and tissues, and the y-axis represents transcripts per million (TPM) values. d) CoNekT comparative co-expression analysis of the four CESA genes indicated in panel b. Nodes represent genes, while solid and dashed edges connect co-expressed and orthologous genes. Coloured shapes indicate the different orthogroups. e) Order tree summarising the duplication events of cell wall-related genes. The tree is based on the gene tree of CESAs and lignin-related genes. Coloured shapes represent the different gene classes.

### Fern-specific evolution of secondary cellulose synthases

To better understand the function of the duplicated CESAs ferns, we first analyzed the gene tree of two CESA clades (Figure 5a, red and blue arrow). Both clades contain genes from *Dicranopteris curranii* (red clade: *Dcu_g31359,* blue clade: *Dcu_g12277*, black arrows), suggesting a duplication in the ancestor of Gleicheniales-II (Figure 5b). To suggest the function of the red and blue clades, we first examined the expression profiles of two representative genes (*Ceric.13G049300.1*) and (*Ceric.09G024100.1*) from *Ceratopteris richardii* (Figure 5b, blue and red solid arrows, respectively). While *Ceric.13G049300.1* showed the highest expression in vegetative fronds, *Ceric.09G024100.1*’s expression was highest in leaf and shoot tips, suggesting different biological processes for the two genes (Figure 5c).

We next compared the co-expression network of *Ceric.13G049300.1* to other ferns using the CoNekT’s Expression Context Conservation (ECC) panel (https://conekt.sbs.ntu.edu.sg/sequence/view/2353618). The fern gene with the most similar expression network was *Stenochlaena palustris Spa_g26805*, which happens to be found in the same clade (Figure 5b, blue solid and double arrow). The networks of *Ceric.13G049300.1 and Spa_g26805* contain orthologs involved in lignocellulose production, such as *CESAs, 4CLs, OMT1s, CYP98A3* (lignin-related *C3’H*), *CSI1* ^104^ and laccases ^105^(Figure 5d). Thus, the genes from the blue clade are likely involved in secondary well wall biosyntesis, indicating that ferns independently evolved this module. Conversely, *Ceric.09G024100.1* and its most similar co-expression ortholog was *Pyrrosia piloselloides Ppi_g22229* (Figure 5b, green solid and double arrow) were co-expressed with genes unrelated to lignocellulose production (e.g., genes similar to monoterpenol-associated *CYP76C2* ^106^)(Figure 5d). This suggests that the second *CESA* clade might be involved in another unknown biological process. Taken together, this indicated that ferns have independently evolved a secondary cell wall module, and further duplicated the CEASs to perform yet unknown functions.

### The evolution of lignocellulose-biosynthesizing genes in land plants

To better understand how the genes involved in lignin and cellulose synthesis have evolved in land plants, we mapped the timing of gene duplications onto the land plant species tree (Figure 5e). Because both *CESA6-like* and linear *CESA* clades both contain bryophytes (Figure 5a), we propose that the ancestor of land plants contained a *CESA6- like* and a *CESA* of linear-type. The ancestor of ferns and seed plants evolved *CESA1/3- like* genes, that further expanded in seed plants into *CESA1* and *3* and secondary *CESAs4,7,8* (Figure 5a). Within ferns, we observed duplications of *CESA1/3-like* in the ancestor of Gleicheniales-II (Figure 5a, duplication 3), *CESA6-like* in Hymenophyllales (duplication 4), and two duplications of the fern-specific CESAs (duplications 1,2) in Gleicheniales-II. We also observed a complete gene set of lignin biosynthetic genes in early-diverging land plants, and evidence of duplication of *4CL* in the ancestor of ferns and seed plants and the ancestor of Hymenophyllales (Figure S19).

Taken together, the ferns show a prolific duplication of *CESA* genes deeply within the fern lineage (Figure 5e), further suggesting that ferns have evolved cell walls with yet unknown features.

## Discussion

Despite ferns’ critical evolutionary position as the sister group, no large-scale studies that investigated their phylogeny, biological pathways and cell walls had been performed. To remedy this, we generated gene expression atlases for 22 ferns and covered ten out of 12 fern orders (Figure 1), allowing us to generate a high-quality species tree that resolved the long-standing relationship between Gleicheniales and Hymenophyllales ^12,107,108^. The tree is supported by outgroups comprising lycophytes, horsetails or seed plants, but not bryophytes. We speculate that the greater phylogenetic distance between bryophytes and the other lineages, combined with reduced single-copy gene dataset obtained from OrthoFinder (Supplemental Methods 1), and degenerated phylogenetic signals in amino acid sequences contributed to the discordance between bryophyte-based and the other outgroups.

The species tree of ferns allowed us to estimate the time of speciation and whole genome duplication events. The WGD analysis revealed that WGD events likely contributed to species diversity (Figure S3), but we observed no correlation between the number of WGDs and genome size. This suggests that alternative evolutionary sources contribute to the exceptional genome size of ferns, and that recent transposon activities and ploidy variation might play a bigger role.

The stem age of early diverging ferns Equisetidae (consisting of Equisetales), with 95% HPD time estimates are in line with the estimate from ^109^ and the oldest unequivocal euphyllophyte fossils ^1^. Polypodiidae was originated in the time range between Lower Carboniferous (323.2 ± 0.4 - 358.9 ± 0.4 mya) and Middle Devonian (382.7 ± 1.6 - 393.3 ± 1.2 mya), with 95% HPD time estimate as 345.09 - 389.61 mya and posterior mean 369.01 mya, which might have first survived the Hangenberg and Kellwasser extinction events before its substantial diversification. The early diverging leptosporangiate fern order Osmundales originated amid Upper to Lower Carboniferous (298.9 ± 0.15 - 358.9 ± 0.4 mya), consistent with ^13^. The aquatic Salviniales, the only extant ferns with heterospory, was originated between Upper Triassic (201.4 ± 0.2 - 237 mya) and Lower Permian (273.01 ± 0.14 - 298.9 ± 0.15 mya), with 95% HPD time estimate as 211.15 - 273.19 mya and posterior mean 241.70 mya, which might correlate with the P-T event after which a vast majority of aquatic environment became empty and the innovative microspores might facilitate their spread and survival. The two most species-diverse suborders, Polypodiineae (i.e., eupolypods I) and Aspleniineae (i.e., eupolypods II) of the Polypodiales were originated amid the Cretaceous (66.0 - 145.0 mya), a relatively warm and ice-free period, with 95% HPD time estimate as 89.75 −144.92 mya and posterior mean 115.93 mya, coincident with the burst of angiosperms in the mid-Cretaceous as highlighted by Darwin ^110^ and the decline of gymnosperms ^111^.

Our gene inventory analysis shows massive gains of genes in the fern lineage (Figure 2a), resulting in ∼50% being fern-specific (Figure 2b). Expression analysis revealed that the fern-specific genes tend to be organ-specific, suggesting their role in fern-specific adaptations. Conversely, older genes are ubiquitously expressed (Figure 2c), which aligns with our previous observation that these genes tend to have basal, essential functions (e.g., photosynthesis, protein synthesis, DNA duplication)^47^. Many of the genes involved in angiosperms’ hormonal and developmental pathways were missing (Table S9), showing that ferns have organised these pathways differently.

Signalling and biosynthetic pathways may significantly vary within land plants, and these pathways tend to expand to support increased anatomical and lifestyle complexity ^112^.Thus, the arguably simpler fern hormonal pathway genes might suggest that these pathways can function in ferns without their angiosperm counterparts. Alternatively, ferns might have evolved equally complex but alternative signalling pathway components that show no sequence similarity to known angiosperm genes. This idea is exemplified by our analysis of flavonoid biosynthesis genes. The lack of these enzymes in ferns–but the presence of flavonoids–indicates that the ‘canonical’ flavonoid pathway is an angiosperm-specific invention and suggests that ferns have either convergently evolved other enzymes with similar functions or use a different pathway to synthesise these compounds. As fern-specific genes are co-expressed with genes involved in development, reproduction and various signalling pathways (Figure 2e), ferns likely have independently expanded these pathways.

The observed high amounts of lignin S units in *P. irregularis* and *S. palustris* (Figure 3ab), and the absence of angiosperm- or lycophyte-specific *F5H* enzymes suggest that the S unit has independently evolved at least four times in the plant lineage: angiosperms, lycophytes, gymnosperms and now ferns ^113^. The re-emergence of S lignin in distantly related plant lineages implies that it may have an essential role in plants’ environmental adaptation, such as improved mechanical properties or herbivore resistance ^113^. While lycophytes have evolved a C3’H-independent pathway by inventing a dual meta-hydroxylase *SmF5H* (Figure 3c, blue pathway)^69^, we observed the presence of *C3’H* genes in ferns (Figure 3d), suggesting that ferns have independently evolved a *F5H* enzyme, and likely follow the biosynthetic route of angiosperms. By combining phylogenetic and co-expression analysis, we propose that the red clade shown in Figure 3h, which contains the highest density of *CYP450s* co-expressed with the lignin biosynthetic genes, comprises the fern *F5H* enzymes. The biosynthetic activity of these genes could be tested by in vitro studies, as done for Selaginella Sm*F5H* ^69^.

Our comparative analysis of cell wall-related genes indicated that ferns and angiosperms contain similar gene sets but that ferns have smaller acetyl and methyl- transferase gene families (Figure 4c). Cell wall composition varies considerably between fern species, corroborating findings in earlier glycan array surveys of ferns ^114^. Surprisingly, we observed a clear mixed-linkage *β*-glucan signal from *O. pendulum*, *Angiopteris sp1*, *A. capillus-veneris* and *S. palustris* (Figure 4d), demonstrating that this unusual polymer is found outside of fern Equisetum ^87,88^. While the AGP epitopes showed weak signals, ferns contain AGPs with special features, such as 3-*O*-methylrhamnose, that are not known in angiosperms ^73,115^.

Surprisingly, we observed a wide-spread occurrence of mixed-linkage glucans outside of Equisetidae (Figure 4d), and we propose two candidate enzyme families that could produce this hemicellulose. First, bryophytes, ferns and Selaginella both contain MLGs ^96^ and linear CesAs (Figure 5a). In the moss *Physcomitrium*, linear CesAs produce arabinoglucan ^116^, and the authors point out that these CESAs are related to an ascomycete MLG synthase and thus represent an early system for MLG-synthesis. A second candidate could be the fern CSLB/G-related clade, as the functions of these genes are currently unknown in ferns and angiosperms. The biosynthetic activity of these enzymes could be tested in vivo, as done for the barley MLG synthase ^117^.

Separate sets of CesAs for primary and secondary cell wall synthesis are a shared feature of spermatophytes ^91^. Whereas tracheids probably evolved once uniting all tracheophytes ^118^, vessels have evolved in angiosperms and independently in *Selaginella, the Gnetales, Equisetum* and other ferns ^119^. Surprisingly, *Selaginella* does not have two sets of CesAs, and we did not observe CesA related to angiosperm secondary cell walls in ferns (Figure 5a), suggesting that their vascular elements result from convergent evolution ^96^. Convergent evolution is supported by a fern-specific CesA clade containing genes involved in lignocellulose biosynthesis (Figure 5). We observed several duplications of the cell wall-related genes within ferns (Figure 5e), which aligns with similar observations in angiosperms and bryophytes ^120,121^. Combined with the presence of a non-canonical sugar 2-O-Methyl-D-glucopyranose observed in Polypodiales (Figure 4a), our data suggest that cell walls underwent independent innovations within ferns. While it is unclear whether 2-*O*-Methyl-D-glucopyranose is biosynthesized by ferns or bacterial or fungal organisms found in the environment, their significant presence in the fern cell walls indicates that they might have a role in fern biology.

We anticipate that the availability of the comprehensive coding sequence and transcriptomic data from ferns - and their availability as a user-friendly CoNekT database (https://conekt.sbs.ntu.edu.sg/)-will be mined to lead to vital insights into the evolution of plant genes and gene families. Implementing fern data into the existing comparative genomic framework will enhance our understanding of the plant tree of life.

## Methods

### Sampling of ferns

22 ferns from 22 families were sampled across Singapore (Table S1). Fern organs were sampled as three biological replicates, where organs were selected to capture the highest variance in the developmental stages and morphological characteristics (Figure S1). Samples were placed into 15 ml falcon tubes and kept in liquid nitrogen and subsequently at −80°C to prevent degradation of RNA.

### RNA isolation and sequencing

After collection, each sample was ground in liquid nitrogen to a fine powder. RNA was extracted from 100 mg of plant material using Spectrum™ Total Plant RNA Kit (Sigma) Protocol A following the manufacturer’s instructions. Quality control of all extracted RNA was carried out by Novogene (Singapore). Each sample was evaluated for its quantity, integrity and purity using agarose electrophoresis and Nanodrop. Library construction was performed by Novogene, and mRNA was enriched from total RNA with oligo(-dT) magnetic beads. The library was then quantified with Qubit and real-time PCR and sequenced using Illumina NovaSeq 6000, with paired-end sequencing of 150 base pairs (bp) per read and a sequencing depth of approximately 60-70 million reads.

### Transcriptome assembly

Low-quality RNA-seq reads were removed, and the remaining reads were trimmed with Fastp (v0.23.2)^122^. Reads were assembled via a curated transcriptome assembly pipeline (Figure S2). Reads were assembled in three biological replicates for each organ, with all organs concatenated and filtered. Each organ consisting of three reads was assembled using SOAPdenovo-Trans (v1.03)^37^ with 10 single *K-mer* (21-39) and Trinity (v2.8.5)^36^ with 25 single *K-mer*. All reads were concatenated into a Trinity-SOAPdenovo assembly and filtered through the Evidential Gene Pipeline (http://arthropods.eugenes.org/EvidentialGene/) using trformat.pl and tr2aacds.pl for removal of any redundant coding sequences. The filtered transcriptome was evaluated using BUSCO (v5.4.3)^123^ and embryophyta as the dataset. All organ assemblies within the sampled fern species were concatenated and filtered again through the EvidentialGene Pipeline, with the output transcriptome being filtered via transcripts per million (TPM) reads using kallisto (v0.50.1)^124^ against each RNA-seq. TPM scores were averaged per organ, and coding sequences with scores < 1 were removed from the assembly. Assembly was then filtered using GC% content, with redundancy coding sequences with less than 40% and more than 60% removed. All transcriptomes were blasted against the NCBI database, and sequences with a percentage identity of more than 70% and an e-value less than e^-10^ were removed from the final assembly. The quality of assembly was determined by BUSCO using the embryophyta dataset (Table S2).

### Construction of *K*S-based age distributions

*K*S-age distributions for all paralogous genes (paranome) of genomes and transcriptomes were constructed by ksrates (v1.1.1)^125^. In brief, the ksrates pipeline entails firstly translating the coding nucleotide sequences into peptide sequences assuming standard genetic code, filtering out sequences whose sequence length is not divisible by 3, containing invalid codons or in-frame stop codon, after which an all-versus-all BLASTP was implemented with *E*-value set as 1 x 10^−10^ in BLASTP (v2.11.0+)^126^ and the resultant subject-query hit table was fed into MCL (v14-137) ^127^ with clustering inflation factor set as 3.0 to delineate paralogous gene families while filtering out gene families whose size is larger than 200, secondly calling the aligner MUSCLE (v3.8.1551)^128^ under default parameter to obtain a multiple sequence alignment (MSA) at the protein level for each paralogous gene family while filtering out sequence pairs whose gap-stripped alignment length was shorter than 100, which was then back-translated into a codon alignment and subsequently fed into the CODEML function within PAML (v4.9j) ^129^ to acquire the maximum likelihood estimate (MLE) of *K*S values under non-pairwise mode using the default control file defined by wgd (v1.1.1) ^130^ and then calling FastTree (v2.1.11)^131^ upon the peptide MSA under default parameter to attain a midpoint-rooted phylogenetic tree of each paralogous gene family for retrieving the weight of each paralogous gene pair with or without outliers, and eventually building the *K*S-age distribution with de-redundancy achieved by node-weighted method after excluding outliers. The collinear gene pairs (anchor pairs) were identified by i-ADHoRe (v3.0.01)^132^ under the default control file defined by wgd, and the weight values for anchor pairs whose corresponding *K*S values were between 0.05 and 20 were recalculated and reassigned while the weight of remaining pairs was set as zero. For orthologous *K*S-age distributions, the process of MCL clustering was superseded as reciprocal best hits (RBH) searching to identify orthologous gene pairs while the weighting process was revoked on that only one-versus-one orthologues were inferred. CD-HIT (v4.8.1)^133^ was applied for the de-redundancy of transcriptome assemblies with the clustering threshold set as 0.99 before *K*S-age analysis.

### Correction of differences of synonymous substitution rates

Synonymous substitution rates were corrected in ksrates. The principle leans on a number of trios of species, including a focal species, a sister species and an outgroup species. The disparate synonymous substitution rate between focal species and sister species since the divergence is represented indeed by the branch-specific contribution of accumulated synonymous substitutions per synonymous site in respective branches. The mode of orthologous *K*S-age distributions inferred from the kernel density estimate (KDE) using Gaussian kernels within the python package scipy was designated as the proxy of the peak *K*S value of each orthologous *K*S-age distribution. 200 iterations of bootstrap with replacements were implemented for each orthologous *K*S-age distribution, and the mean along with standard deviations (STD) of mode across the replicates was determined as the final peak *K*S value representing divergence distance and its associated STD. The original accumulated synonymous substitutions per synonymous site of focal species- sister species pair consisting of the branch-specific contribution of both species since diversification was transformed into two times the branch-specific contribution of focal species with the prop of outgroup species to resemble the timescale of focal species. The mean of rescaled peak *K*S values of focal species-sister species pair against various outgroup species was taken as consensus-adjusted peak *K*S value. The maximum number of trios was set as 20. The species tree inferred by ASTRAL-Pro2 using seed plants as outgroup species were adopted in ksrates.

### Construction of orthologous families and single-copy gene trees

Orthofinder (v2.5.4)^134^ was performed upon the protein sequences of 107 ferns and outgroup species with an inflation factor set as 3 to delineate the orthologous families. No single-copy gene families were identified by Orthofinder, probably because of the universal and unique gene duplication and loss scenario across species and gene isoforms ^135^. To recover reliable and adequate single-copy gene families, we constructed mostly single-copy gene families ^51^, wherein most species were in single-copy while the remaining species had no more than four copies which were assumed to be transcript variants of the same gene, by retaining the longest copy, if applied, of each species. We referred to the mostly single-copy gene families as single-copy gene families thereafter. In total, 140, 112, 107 and 57 single-copy gene families were constructed from dataset 107 ferns, 107 ferns plus seed plants, 107 ferns plus seed plants and lycophytes, 107 ferns plus seed plants, lycophytes and bryophytes, respectively. MAFFT (v7.475)^136^ was performed to obtain a peptide multiple sequence alignment (MSA) for each single-copy gene family with the parameter “–auto”. Trimal (v1.4.1)^137^ was then performed to trim the MSA and back-translate it into a codon-level nucleotide MSA with parameter “- automated1”. IQ-TREE (v1.6.12)^138^ was implemented on each codon-level nucleotide MSA wherein ModelFinder ^139^ was called to find the best-fit codon substitution model in terms of Bayesian Information Criterion (BIC) upon which a maximum likelihood (ML) gene tree was inferred and assigned with bootstrap support values from 1000 ultrafast ^140^ bootstrap replicates with parameter “-bnni” to further optimize each bootstrap iteration through a hill-climbing nearest neighbor interchange (NNI) search based directly on the corresponding bootstrap alignment to avoid severe model violations. The same process was further applied to the 108 ferns dataset including *Marsilea vestita*, in which a total of 136, 108, 103 and 55 single-copy gene families were reconstructed from datasets varied in outgroups with both nucleotide and peptide molecules.

### Species tree inference

Three methods, ASTRAL-Pro2 ^41^, STAG ^42^, and a concatenated-based method were implemented to infer the species tree. The acquired individual ML single-copy gene trees and gene name-species name map files were imported into ASTRAL-Pro2 and STAG under default parameters to estimate a consensus species tree with support values for each bipartition denoting local posterior probabilities (localPP) and the proportion of individual estimates of the species tree that contain that bipartition, respectively. For the concatenated-based method, the individual codon-level nucleotide MSA of single-copy gene families were concatenated and then fed into IQ-TREE to infer a ML super- gene tree as above. *Dicranopteris curranii*, *Dicranopteris pedata*, *Diplopterygium laevissimum*, *Diplopterygium glaucum* and *Sticherus truncatus* in the Gleicheniaceae were named as Gleicheniales−II clade while *Cheiropleuria integrifolia*, *Dipteris conjugata* and *Dipteris lobbiana* in the Dipteridaceae were named as Gleicheniales−I clade p. In total, (140), (112), (107) and (57) single-copy gene families were constructed from the (107 ferns dataset), (107 ferns plus seed plants), (107 ferns plus seed plants and Lycopods), (107 ferns plus seed plants, Lycopods and Bryophytes), respectively. The 108 ferns dataset including *Marsilea vestita*, were with 136, 108, 103 and 55 single-copy gene families, respectively.

### Estimation of absolute divergence time

Mcmctree (v4.9j)^129^ was implemented upon the concatenated peptide MSA of single-copy gene families of 108 ferns dataset with Equisetales as outgroup to infer the absolute divergence time for each bipartition. The independent rates model, which assumes a log-normal distribution of evolutionary rates across branches, was selected and 18 fossil calibrations of soft constraint from ^141^ were adopted to refine the divergence time of internal nodes, as summarised in Table S2. Fossils calibrating clades within Gleicheniaceae or Hymenophyllaceae were avoided for their indefinite phylogenetic location. LG amino acid substitution matrix was selected and a gamma model of rate variation was assumed with alpha as 0.5 and 5 categories in discrete gamma. Parameters controlling the birth-death process were set as 1, 1, 0.1 to generate uniform age priors on nodes that didn’t have a fossil calibration. Gamma priors for the transition/transversion rate ratio and shape parameters for variable rates among sites were set as 6 2 and 1 1. A Dirichlet-gamma prior was set upon the mean rate across loci and the variance in logarithm as 2 20 1 and 1 10 1. The first 2000 iterations were discarded as burn-in and then 20,000,000 iterations were performed with sampling per 1000 iterations. The effective sample size (ESS) of all parameters was larger than 200, suggesting adequate sampling and convergence.

### Phylogenomic analysis of gene tree - species tree reconciliation

To estimate the retention rate and interrogate hypothetical WGDs over competing scenarios in different clades, we implemented 4 categories of statistical gene tree - species tree reconciliation analysis using Whale (v.2.0.3)^142^, as shown in Figure SM9-12. Firstly, orthogroups of each category of species were inferred by Orthofinder (v2.5.4) ^134^ with an inflation factor as 3. Gene families were filtered to assure at least one gene from each descendant present at the root and to avoid large gene family size(which contains noise and causes computational downshift) via “orthofilter.py” (https://github.com/arzwa/Whale.jl)1000 gene families were randomly selected as subsequent inputs. PRANK (v.150803)^143^ was utilised to obtain a MSA for each gene family and MrBayes (v.3.2.6)^144^ was then applied to infer the posterior distributions of gene trees under the LG + GAMMA model, with iterations set as 110,000 and sample frequency as 10 to get in total 11,000 posterior samples. ALEobserve ^145^ was subsequently performed on the tree samples to construct the conditional clade distribution with a burn-in of 1000. Two gene family evolution models, the relaxed branch-specific model and the critical branch-specific model, were applied as in a previous study ^45^ to estimate the retention rates of hypothetical WGDs for each category, as shown in Figure SM9-12. Hypothetical WGDs with retention rates higher than 0.05 were regarded as supported WGDs, considering the incompleteness of transcriptome assemblies and the stochasticity of sampled gene families.

### Absolute dating of WGDs

Phylogenetic dating of AZOL and CERA WGD proceeded as follows. Firstly, an orthogroup comprising orthologues from 8 other species and anchor pair which was assumed to be retained from the corresponding WGD was constructed per anchor pair by searching the reciprocal best hits (RBH) between the anchor pair and the transcriptomes or genomes of other species by Diamond (v2.0.5.143)^146^ under default parameters (Figure SM14). *K*S range 0.36 - 2.00 was confined for the age of anchor pairs to be adopted in terms of densest aggregation of duplicates and avoidance towards saturation for AZOL WGD and *K*S range 0.41 - 2.0 was bounded for CERA WGD. Secondly, the peptide sequences of each individual orthogroup were aligned by MAFFT (v7.475)^136^ under default parameters and then concatenated as a single peptide MSA. The numbers of concatenated orthogroups were 45 and 14 for AZOL and CERA WGD, respectively. The adopted fossil calibrations followed Table S2 at corresponding phylogenetic locations while the boundaries of root for AZOL WGD were set as minimum bound 168 mya based on the minimum bound of fossil calibration “Stem Lygodiaceae” and safe maximum bound 345 mya as the fossil calibration “Stem Osmundaceae”, as shown in (Figure SM14). Mcmctree (v4.9j)^129^ was implemented for the Bayesian molecular dating for each WGD with the parameters same as above. The ESS of all parameters was larger than 200, indicating adequate sampling and convergence. The posterior distribution of time estimate for the node joining the anchor pair was retrieved and the 95% HPD, posterior mean, median and mode were adopted to characterise the age of WGD, as shown in (Figure SM13).

### Identifying organ-specific genes

Organ-specific genes were isolated from each transcriptome via specificity measure (SPM) values ^47^. For each gene, we calculated the average TPM values in each organ. Following that, the SPM value of a gene in an organ was calculated by dividing the average TPM in the organ by the sum of the average TPM values of all organs. The SPM value ranges from 0 (gene not expressed in organ) to 1 (gene fully organ-specific). To identify organ-specific genes for each organ, we first identified an SPM value threshold above which the top 5-11% of SPM values were found (Figure S5). These top values varied across the species sampled, depending on the number of organ-specific genes identified. If the SPM value of a gene in an organ was equal or greater than the threshold value, the gene was identified as organ-specific within said organ. Organ-specific genes were then plotted in a heat map to show their distributions (Figure S6).

### Functional annotation of genes

Assembled sequences of 22 fern species, including four ferns with genomes available online (*A. spinulosa, A. filiculoides, C. richardii* and *S. cucullata*) were annotated using the online tool Mercator4 v.2.0 ^147^. We visualised the Mercator4 annotation using a heatmap, showing the distribution of Mapman Bins across sampled fern species (Figure S9).

### Assignment of orthogroups to phylostrata

Using the coding sequences of the transcriptomes, we constructed orthologous gene groups (Orthogroups) with Orthofinder (v.2.5.4)^134^. Respective outputs for orthogroups were used for further analysis. By utilising the theoretical evolutionary line produced by the phylogenomic analysis of gene trees, phylostratic nodes were assigned to orthogroups based on plant lineages. This analysis spanned a total of 47 species across the plant kingdom, and assigned nodes ranging from node 1 (most ancient, ancestor of Archaeplastida) to node 13 (ancestor of Polypodiales). Nodes were assigned based on the fern species tree (Figure 1c), as well as known phylogenetic analyses of early plants ^148^. The nodes are: node 1 (ancestor of Archaeplastida), node 2 (ancestor of green plants), node 3 (ancestor of land plants), node 4 (ancestor of vascular plants), node 5 (ancestor of ferns and seed plants), and node 6-13 (various fern orders). Specifically, Equisetales is designated as node 6, Psilotales and Ophioglossales as node 7, Marattiales as node 8, Hymenophyllales and Gleicheniales-II as node 9, Schizaeales as node 10, Salviniales as node 11, Cyatheales as node 12, and Polypodiales as node 13. Species-specific gene families were characterised by gene families consisting of only one species, and hence, not assigned to nodes. In cases where nodes encompass multiple species, such as node 4, orthogroups containing only one node assignment (e.g., those with genes solely from Ophioglossales and Psilotales) were not designated to specific nodes.

### Orthogroup gain loss analysis

Gain and loss of orthogroups were determined by the presence of an oldest clade member in a particular node. Potential contamination by non-fern sequences due to the nature of transcriptome assembly was filtered out at this stage by checking for the presence of at least half of the expected clades in each node. For basal nodes (nodes 1 to 4), the clades used were ’Glaucophytes’, ’Chlorophytes’, ’Bryophytes’, ’Lycophytes’, ‘Tracheophyta’ and ‘Spermatophyta’). Nodes were defined as lost based on the clade that they last appeared in.

### Identification of orthogroup expression profiles

Analysis of the expression profiles at phylostrata level was performed as in ^47^, by classifying orthogroups into ‘organ-specific’, ‘ubiquitous’ or ‘not conserved’. Organ-specific orthogroups are orthogroups containing organ-specific genes and were subclassified according to the organ (leaf-, meristem-, crozier-, root meristem-, male-, spore-, rhizome-, root-specific). Orthogroups that are expressed in different organs for each species - that is, that do not show an ‘organ-specific’ expression profile in different species - were labelled as ubiquitous. Orthogroups that had different organ-specific expression profiles in different species (orthogroups containing root-specific genes for *Alsophila latebrosa* and leaf-specific genes for *Equisetum hyemale*) were labelled as not conserved. Only orthogroups that fulfilled the following criteria were identified as organ-specific: (1) Contained at least two species with transcriptome data within each orthogroup. (2) >50% of the genes within the orthogroup supported the expression profile and (3) ≥50% of the species present in the node supported the expression profile.

### Structural analysis of fern-specific orthogroups

25 orthogroups containing at least 10 fern species with protein sequence representatives from sequenced genomes were used to check for sequence similarity by NCBI BLASTp restricted to Viridiplantae (E-value < 1e-10, Query cover > 50%), prediction of structure (alphafold 3 -server)^149^, structure similarity search using DALI (all PDB, Z score > 8, lali > 0.5 of residues in the protein)^150^ and foldseek ^151^. The cif outputs from alphafold 3 were converted to pdb format for input to DALI and foldseek using UCSF ChimeraX version: 1.8. The sequenced representatives were selected based on the highest similarity to the consensus sequence, which was derived from the multiple sequence alignment generated using Seaview v5.0.5 (-align -align_algo 1 -output_format clustal -o) using the muscle algorithm ^152^.

### Constructing co-expression networks and addition of the ferns to the CoNekT database

Co-expression networks were calculated using the CoNekT framework ^67^, and were also used to update the existing database, available at (https://conekt.sbs.ntu.edu.sg/).

### CASA lignin quantification

Methods used for solvent extraction and determination of lignin content by CASA lignin method closely followed the protocol outlined in ^64^. Species organs that were sampled were ground, with solvent extraction in 80% ethanol for woody samples. For non-woody samples, extraction with water using sonication was done first to remove proteins and other water-soluble components. A cysteine stock solution (0.1 g/mL) in 72% sulfuric acid (SA) was prepared by dissolving 10 g L-cysteine in 100 ml SA. 5-10 mg of the solvent extract was placed in a glass vial, where 1.0 mL of prepared stock solution was added, sealed with a Telfon-lined screw cap and stirred at 24 °C (room temperature) via a magnetic stir bar (400 rpm) for 60 mins until the biomass was completely dissolved. The dissolving temperature was decreased to 24 °C to identify a milder condition, allowing convenient operation and minimising interference from carbohydrates. The solution was diluted with deionized water to a volume of 50 or 100mL in a volumetric flask, depending on the lignin content and biomass weight used. Absorbance of the diluted solution was measured at 283 nm (A283) in a 1 cm quartz cell using a UV spectrophotometer against a blank solution (1 mL stock solution diluted to corresponding volume).

### Thioacidolysis

This method is adapted from ^66^. Briefly, 10 mg of alcohol-insoluble cell wall residues were incubated in 3 mL of dioxane with ethanethiol (10%), BF3 etherate (2.5%) containing 0.1% of heneicosane C21 diluted in CH2Cl2 at 100 °C during 4 hr. Three ml of NaHCO3 (0.2 M) were added after cooling and mixed prior to the addition of 0.1 mL of HCl (6 M). The tubes were vortexed after addition of 3 mL of dichloromethane and the lower organic phase collected in a new tube before concentration under nitrogen atmosphere to approximately 0.5 ml. Then, 10 μL of the mixture was trimethylsilylated (TMS) with 100 μL of N,O- bis(trimethylsilyl) trifluoroacetamide and 10 μL of ACS- grade pyridine. The trimethylsilylated samples were injected (1 μL) onto an Agilent 5973 Gas Chromatography–Mass Spectrometry system. Specific ion chromatograms reconstructed at m/z 239, 269 and 299 were used to quantify H, G and S lignin monomers respectively and compared to the internal standard at m/z 57, 71, 85.

### Neutral sugar content analysis

This method is adapted from ^66^. Neutral monosaccharide sugar content was determined by gas chromatography after acid hydrolysis and conversion of monomers into alditol acetates as described in Hoebler et al., 1989, and Blakeney et al., 1983. Gas chromatography was performed on a DB 225 capillary column (J&W Scientific, Folsorn, CA, USA; temperature 205 °C, carrier gas H2). Calibration was made with standard sugar solution and inositol as internal standard.

### Synthesis of methylated sugars (Figure S14)

Please see supplemental methods. The structures of compounds were ascertained by NMR spectroscopy and were in agreement with reported data.

### Constructing phylogenetic trees for genes controlling primary and secondary cell wall formation

Genes controlling primary and secondary cell wall formation were identified from a previous study of *A. thaliana* ^120^. Using the known genes as reference, we utilised OrthoFinder (v.2.5.4)^134^ with 49 species within Table S9. Genes that were grouped in the same orthogroups with the reference genes were aligned via MUSCLE (v.5.1)^128^ and analysed via IQTree (v.1.6.12)^138^ to construct trees with bootstrap values of 100 (for gene families with more than 600 genes) and 1000 (for those less than 600 genes). Trees were then visualised using MEGAX (v.1.0)^153^, with bootstrap values less than 80% being condensed. Ancestor-specific duplication events were inferred using trees generated using previous phylostratigraphic analysis, where branches containing common earliest ancestors were deemed as such.

### Visualisation of genes controlling primary and secondary cell wall formation

Annotations of fern genes were based on the phylogenetic trees in the methods mentioned above, with genes annotated where reference genes from *A. thaliana* were found. Coexpression coefficients of each gene within 26 fern species were calculated using Pearson’s correlation coefficient (PCC) and transformed into Highest Reciprocal Rank (HRR)^154^ and genes of interest (GOIs) were isolated. Coexpressed GOIs were visualised using Cytoscape (v 3.10.1) (https://cytoscape.org/).

### Analysis of CAZymes and HRGPs

Protein files from coding sequences of 39 plant species were submitted to the dbCAN2 pipeline ^155^, which annotates CAZymes using three tools (HMMer against the CAZyme domain database; DIAMOND for BLASTP against the CAZyme database; dbCAN-sub for HMMER detection of putative CAZy substrates). A majority vote was used and all annotations were supported by two or more tools, which were used to filter for relevant CAZy families. CAZy families, as well as respective functionally described enzymes, were aligned using MAFFT ^136^ (preferably L-INS-i; if the sequence dataset was too big the automatic mode was used). Sequence alignments were submitted to FastTree in default mode ^131^, and homologs of functionally described enzymes were filtered using iTOL ^156^. For DUF families (DUF579, DUF231) and selected other enzymes (CGR2-3, BS1, DARX1, P4H, QUA2 and QUA3), BLASTP was used with the described family members against the protein files as a database. E-value of e⁻⁷ was used, with the rest followed as described above.

Annotation of hydroxyproline-rich glycoproteins (HRGPs) was performed by using the workflow described in ^73^. The protein sequences were filtered first for the presence of N-terminal signal peptides and then classified into 24 classes based on the presence of distinct amino acid motifs and biases (as outlined in ^157^).

### Comprehensive Microarray Polymer Profiling (CoMPP)

The CoMPP analysis was performed according to the method reported by ^86^. Each sample was weighed out in triplicate of 10 mg AIR. The samples were sequentially treated with 300 μL 50 mM trans-1,2-diaminocyclohexane-N,N,N′,N′-tetraacetic acid (CDTA) pH 7.5, followed by extraction with 300 μL 4 M NaOH containing 0.1% (v/v) NaBH4. Each extraction step was carried out for 2 h in a TissueLyser II (Qiagen AB, Sollentuna, Sweden) at 6 s-1 at room temperature. After each extraction, samples were centrifuged for 10 minutes at 4000 rpm, and the supernatant was collected. The samples were added to a 384 well plate and four dilution points were prepared for each sample, then two technical replicates printed on nitrocellulose using an ArrayJet Marathon printer (ArrayJet, Roslin, UK).

Separate arrays for each probe were first blocked with 5% (w/v) low-fat milk powder solution in phosphate-buffered saline (MP/PBS), then probed with a set of specific primary monoclonal antibodies (LM, JIM and MAC 207; Plant Probes, Leeds university, CCRC; Complex Carbohydrate Research Center, University of Georgia, BS-400; Biosupplies Australia and INRA-RU donated from Marie Christine Ralet, INRA, France) (Table S14) for 2 h. After three washes with PBS, the arrays were incubated with 1:5000 solutions of either anti-mouse or anti-rat secondary antibodies (depending on the source of the primary antibody) conjugated with alkaline phosphatase for another 2 h. Following three washes with PBS the array had a final wash in Milli-Q water. Arrays were developed with a 5-bromo-4-chloro-3-indolyl-phosphate (BCIP)/nitro-blue tetrazolium chloride (NBT) substrate andscanned using a flatbed scanner (CanoScan 9000 Mark II; Canon, Søborg, Denmark) at 2400 dpi converting the dots to grayscale. The calculated intensity of the signal was quantified using the microarray analysis software ProScanArray Express (PerkinElmer, Waltham, Massachusetts, USA). The relative intensity values were normalised to a scale from 0 to 100 and transformed into a heatmap.

### Enrichment and depletion of biological processes in fern-specific genes and neighbourhood

Fern-specific genes were defined as genes within orthogroups located in nodes 6-13 (Figure 2a), excluding those orthogroups that contained only genes from one species. Neighbours of fern-specific genes were defined as genes that are co-expressed with a fern-specific gene and are not fern-specific genes themselves. The functional annotations of genes were retrieved (first-level Mapman bins) and subjected to enrichment and depletion analysis against a background of genes assigned to orthogroups. The analysis was performed for each fern using a hypergeometric test and adjusted for multiple testing via Benjamini-Hoechberg correction (q < 0.05)^158^. The overall trend of enrichment or depletion of biological processes across fern species was derived by subtracting the number of depleted Mapman bins across all species from the number of enriched bins.

#### Data availability

The raw sequencing data is available at E-MTAB-13848, while the CDS and protein sequences are found at https://doi.org/10.6084/m9.figshare.26347330. The co- expression networks are available at https://conekt.sbs.ntu.edu.sg/species/.

## Acknowledgements

Jonatan Fangel is acknowledged for contributions to Figure 4, panel e and Sylviane Daniel for help with lignin biochemistry. We thank D. Maizels (http://www.scientific-art.com/) for the illustrations in Figure 1. MM thanks Singaporean Ministry of Education grant MOE-T2EP30122-0017 ‘A Kingdom-wide Study of Genes, Pathways, Metabolites, and Organs of Plants’ for funding. LP and BC (project-number 440046237) and JdV (project-number 440231723; VR 132/4-2) are grateful for funding within the framework of MAdLand (http://madland.science), priority programme 2237 of the German Research Foundation (DFG). JdV further thanks the European Research Council for funding under the European Union’s Horizon 2020 research and innovation programme (Grant Agreement No. 852725; ERC-StG “TerreStriAL”). SdV acknowledges funding by the Lower Saxony Ministry of Science and Culture (Niedersachsen Vorab initiative) and the DFG project 515101361. We thank Luc Saulnier for discussion and help with the identification of the unknown sugar. We also thank Elizabeth Haswell (https://elizabethhaswell.carrd.co) for her help with proofreading the manuscript.

## Supplementary Methods

**Supplementary Methods 1: Inferences of species tree, whole genome duplications, analysis of salicylic acid- and jasmonic acid-mediated signalling in ferns, chemical synthesis of methylated sugars.**

## Supplementary tables

**Table S1. 22 species of Tracheophyta from 22 different families.** Samples were collected from various locations in Singapore, with each species having multiple organs harvested.

**Table S2. Sequencing statistics.** The columns contain descriptions of the 415 sample names, including the species, organs, organ types, and data statistics.

**Table S3. Transcriptome assembly statistics for the 22 ferns.** BUSCO value, MapMan annotation percentage, number of transcripts, GC% content, N50, BUSCO scores and percentage of genes annotated by MapMan are shown.

**Table S4 Clade, order, family, species and source of data of the 108 ferns used in this study.**

**Table S5. Fossil calibrations of soft constraint adopted in this study.**

**Table S6. The 95% HPD and posterior mean age estimates (mya) for the origin of each major clade**

**Table S7. Phylostratigraphic assignments of orthogroups to nodes**. The table shows the orthogroups, clades which are present in the orthogroup and the node where the orthogroup appeared in.

**Table S8. Gene-Organ Specificity.** The table shows the differen species (given by mnemonic), SPM value in a given sample, the number of genes in an organ and the gene ids speciffically expressed in the ogan.

**Table S9. Missing/Present Mapman Bins across 39 species, comprising of Glaucophytes, Chlorophytes, Bryophytes, Lycophytes and Ferns.**

**Table S10. The percentage of annotated clades by MapMan bins.** The species comprising the clades are indicated in column B.

**Table S11. CASA lignin content and thioacidolysis analysis, showing content of H, G and S units.**

**Table S12. BLAST scores of Arabidopsis and Selaginella lignin-related genes against 39 species contained in conekt.sbs.ntu.edu.sg.**

**Table S13. Neutral Sugar Analysis.** The different species and their organs are shown in rows. The columns indicate the abundance of sugars and the standard deviation.

**Table S14. Antibodies, their immunogens and declared specificities and the references where the antibodies were generated/described.**

**Table S15 CoMPP profiles of the 102 cell wall extracts probed with 48 antibodies.** The used solvent and antibodies are shown in columns, the species, organs are shown in rows.

**Supplementary Data 1. Co-expression networks of lignin-related genes, cellulose synthases and cell wall-related transcription factors.** The file should be opened in cytoscape.

---

**Figure S1.**
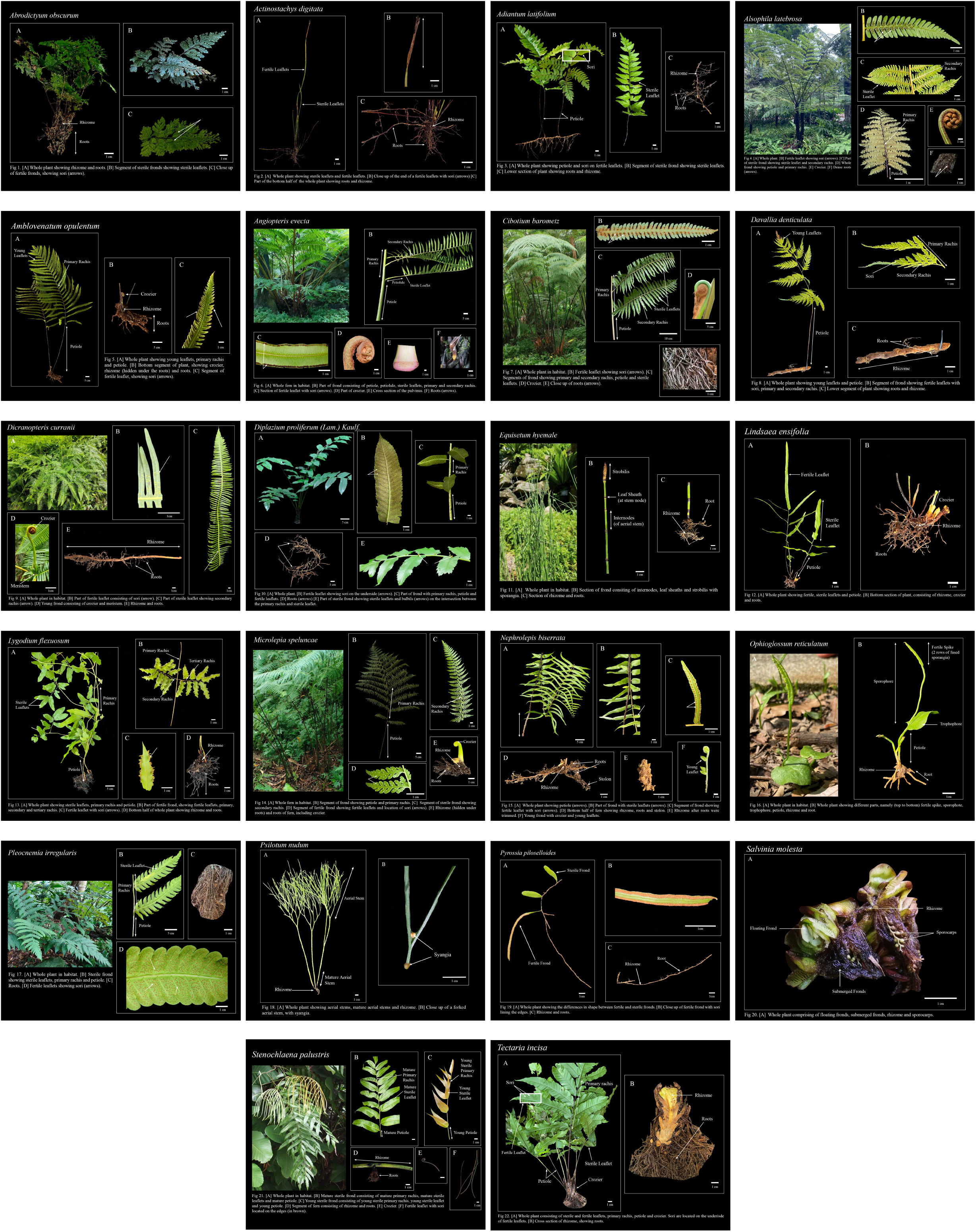
Pictures of the 22 ferns and their sampled organs.

**Figure S2.**
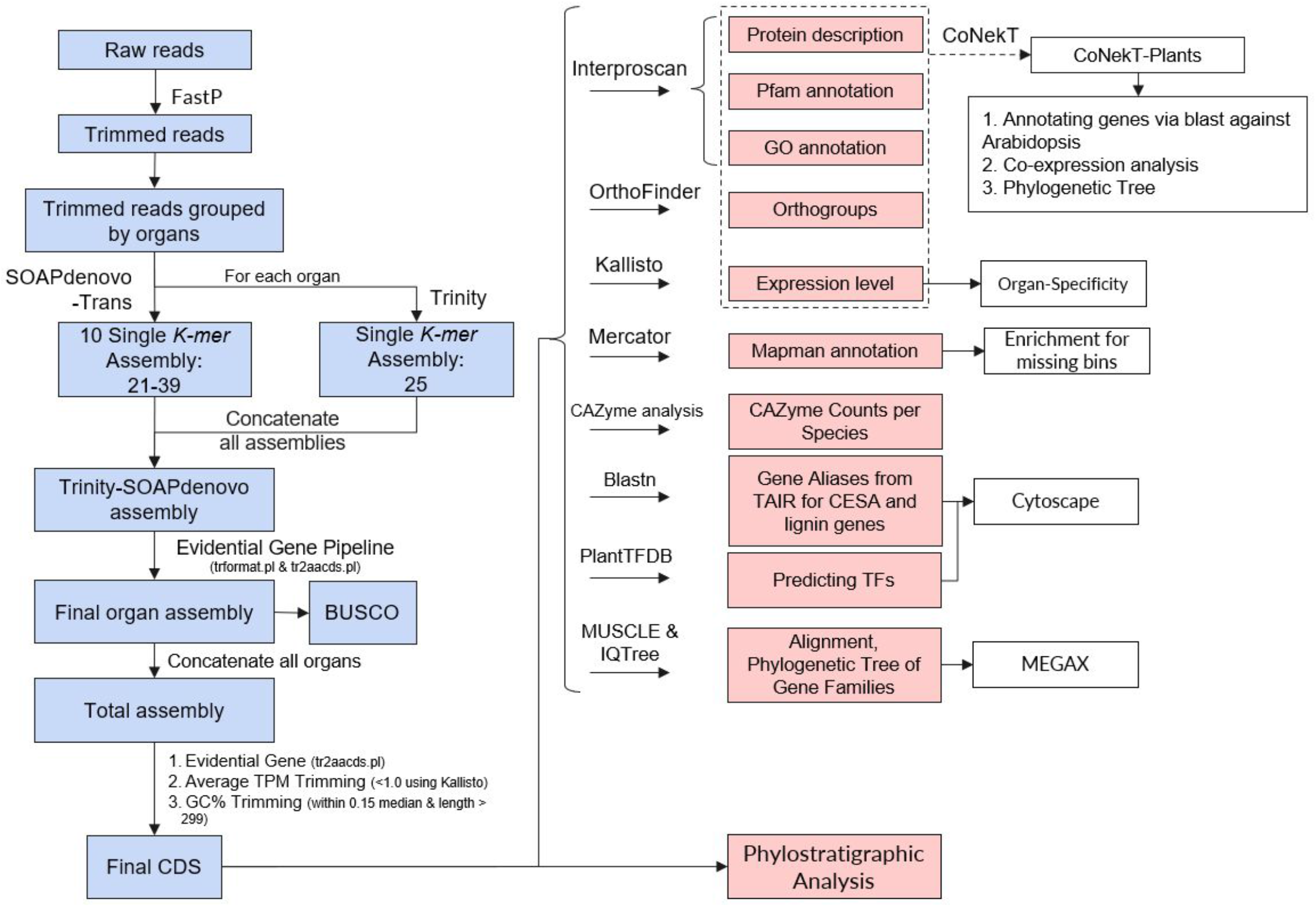
Transcriptome assembly (blue boxes) and subsequent analyses (red boxes).

**Figure S3.**
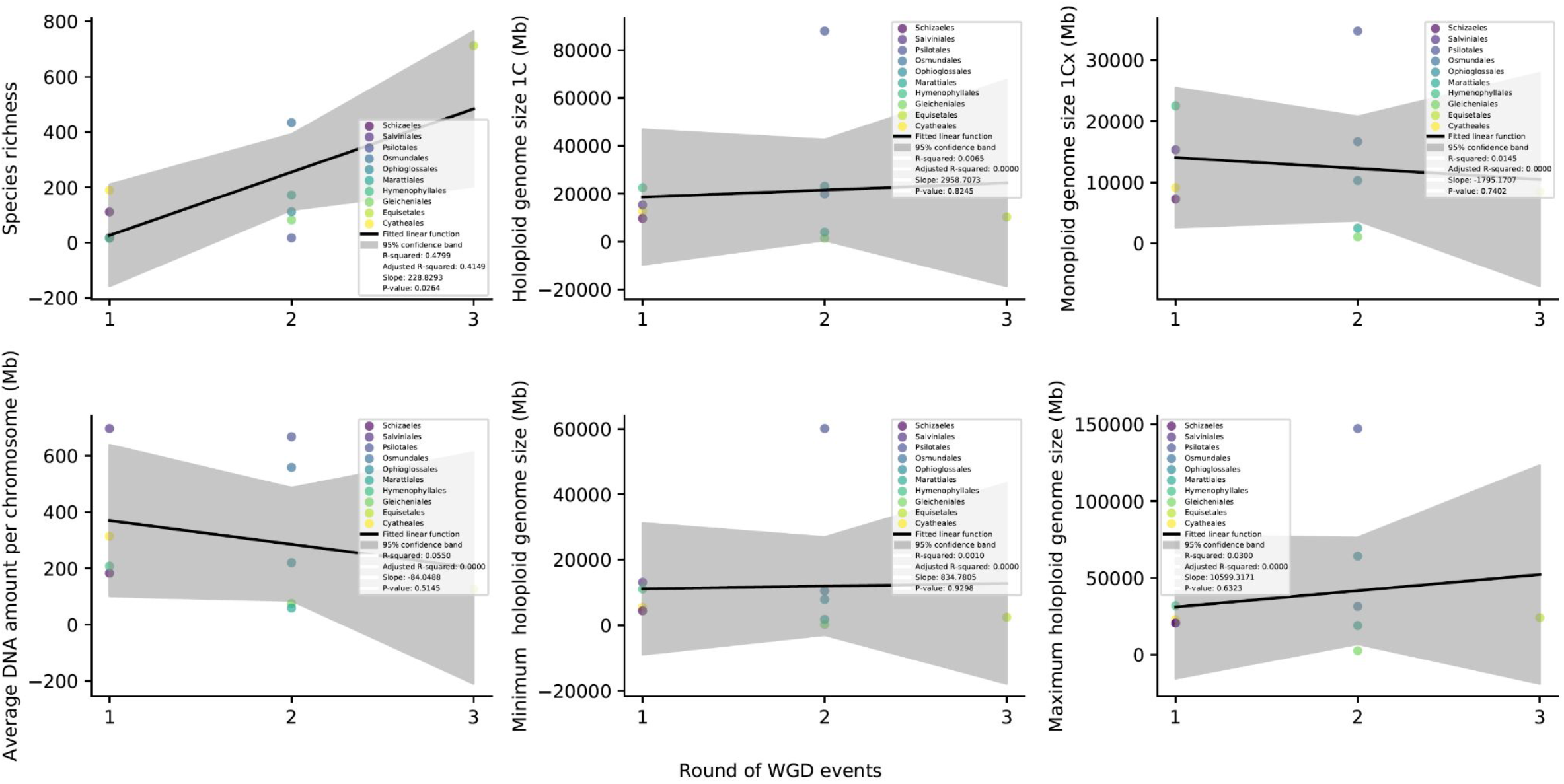
Genomic properties of ferns in relation to whole genome duplication events. The plots show the correlation between WGD events (x-axis) and species richness (the number of species within a lineage), holoploid genome size (total DNA content), monoploid genome size (DNA content of a single set of chromosomes) and others.

**Figure S4.**
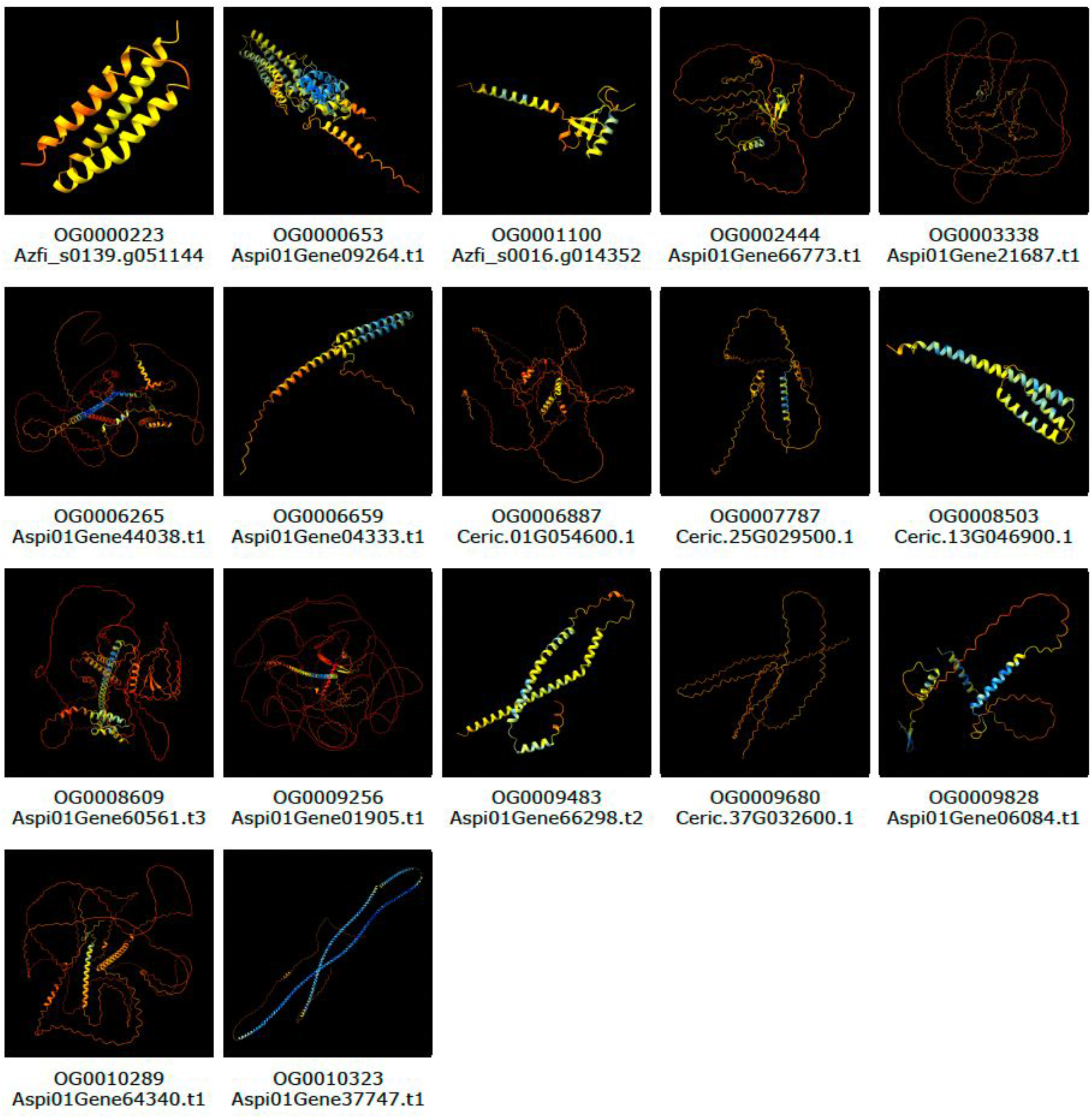
AlphaFold3-derived structures of the 17 fern-specific proteins. The colors indicate confidence scores of the structures.

**Figure S5.**
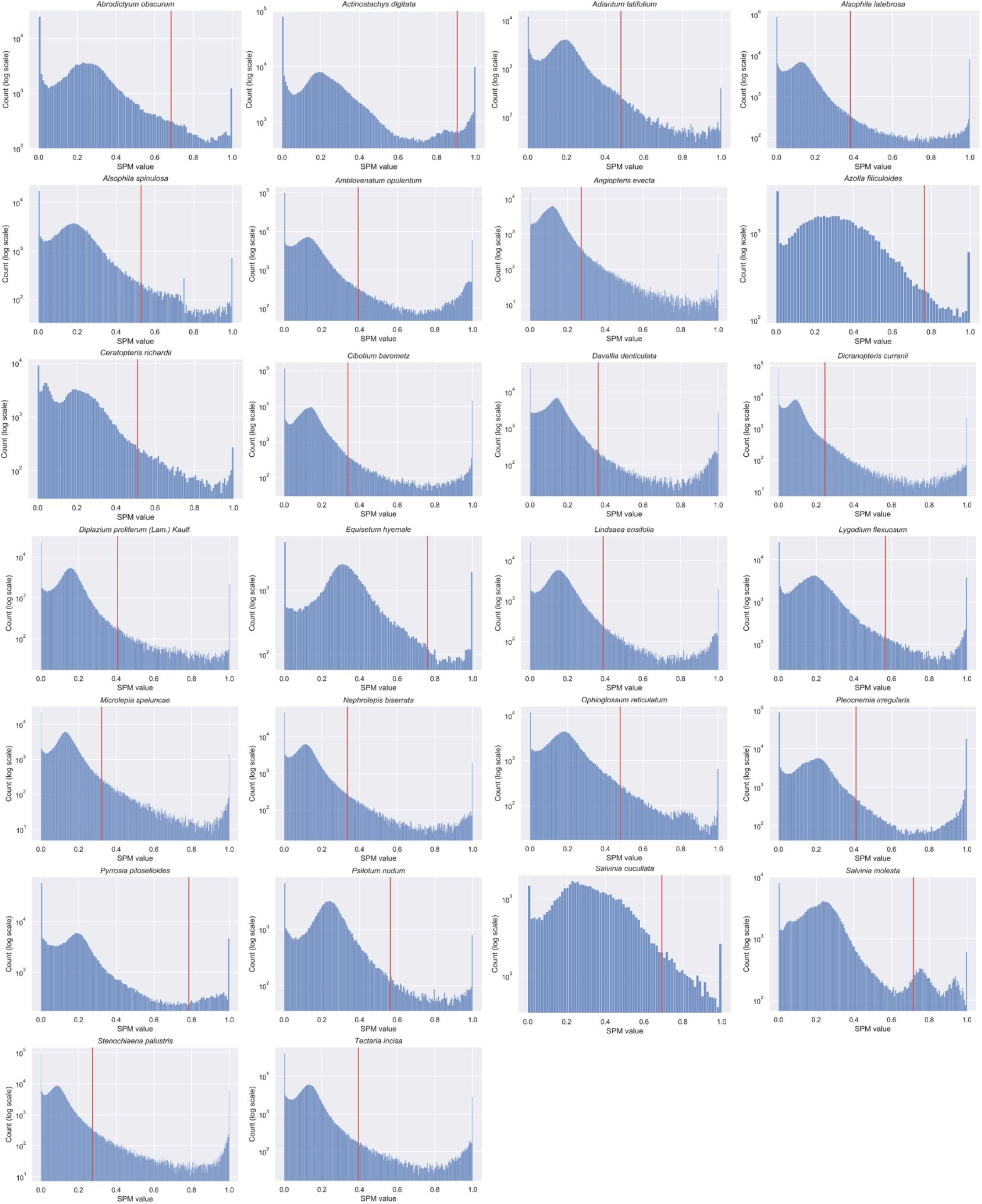
Number of genes (y-axis) with a given SPM value (x-axis). The SPM value cutoff is indicated by the red line.

**Figure S6.**
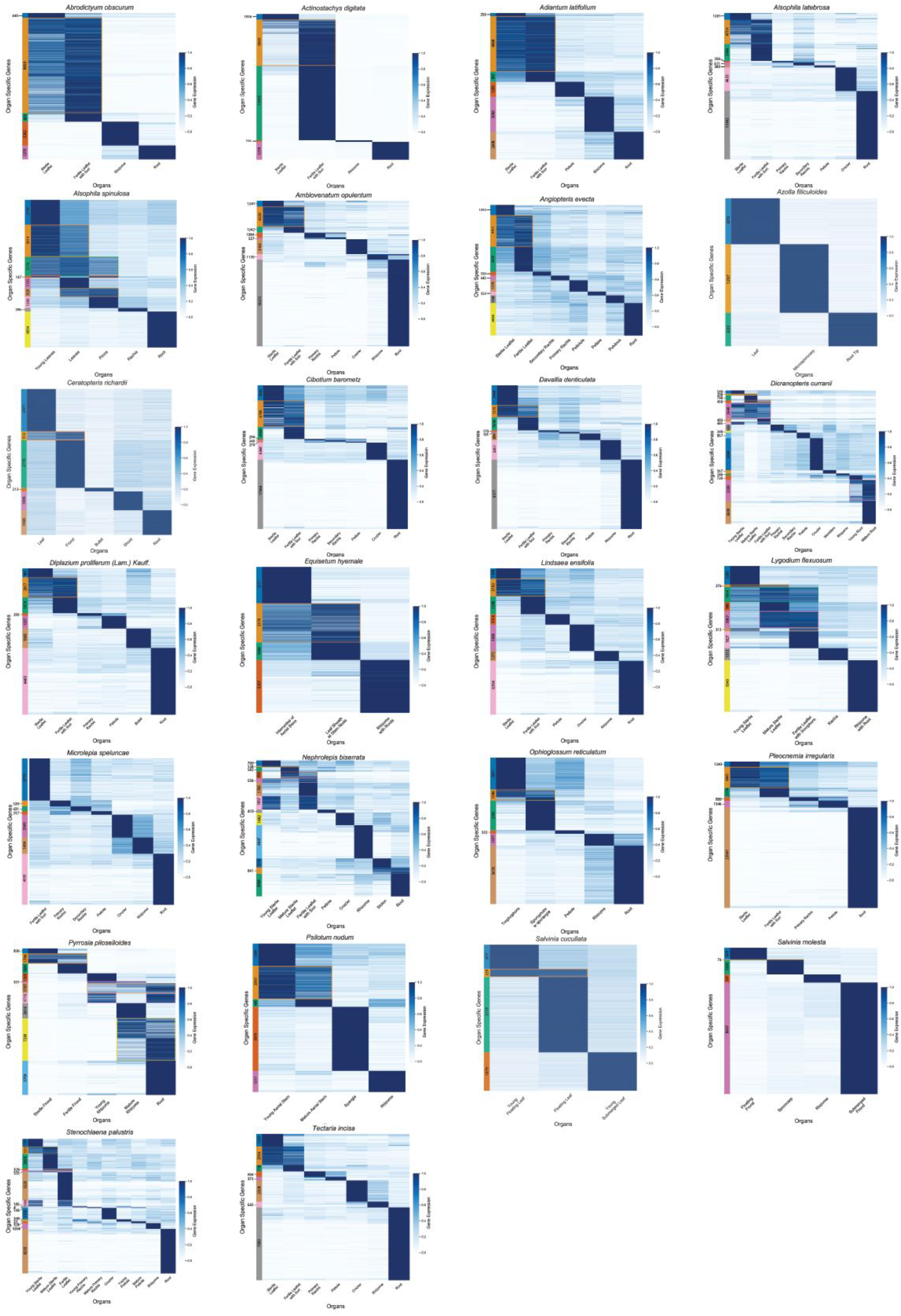
Gene expression profiles of organ-specific genes. Each gene’s expression has been scaled to range from 0 to 1.

**Figure S7.**
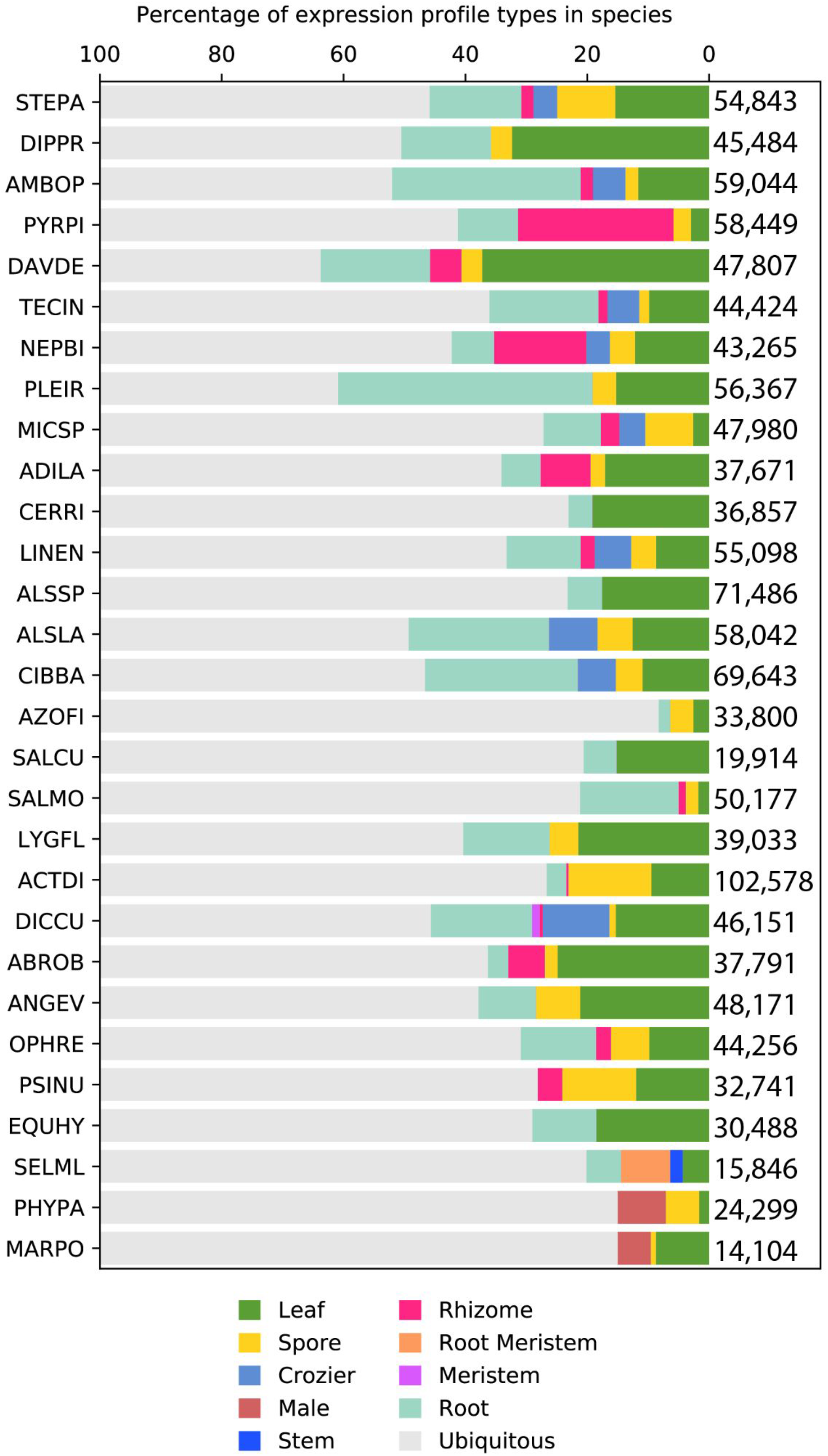
Expression profiles for species-specific genes.

**Figure S8.**
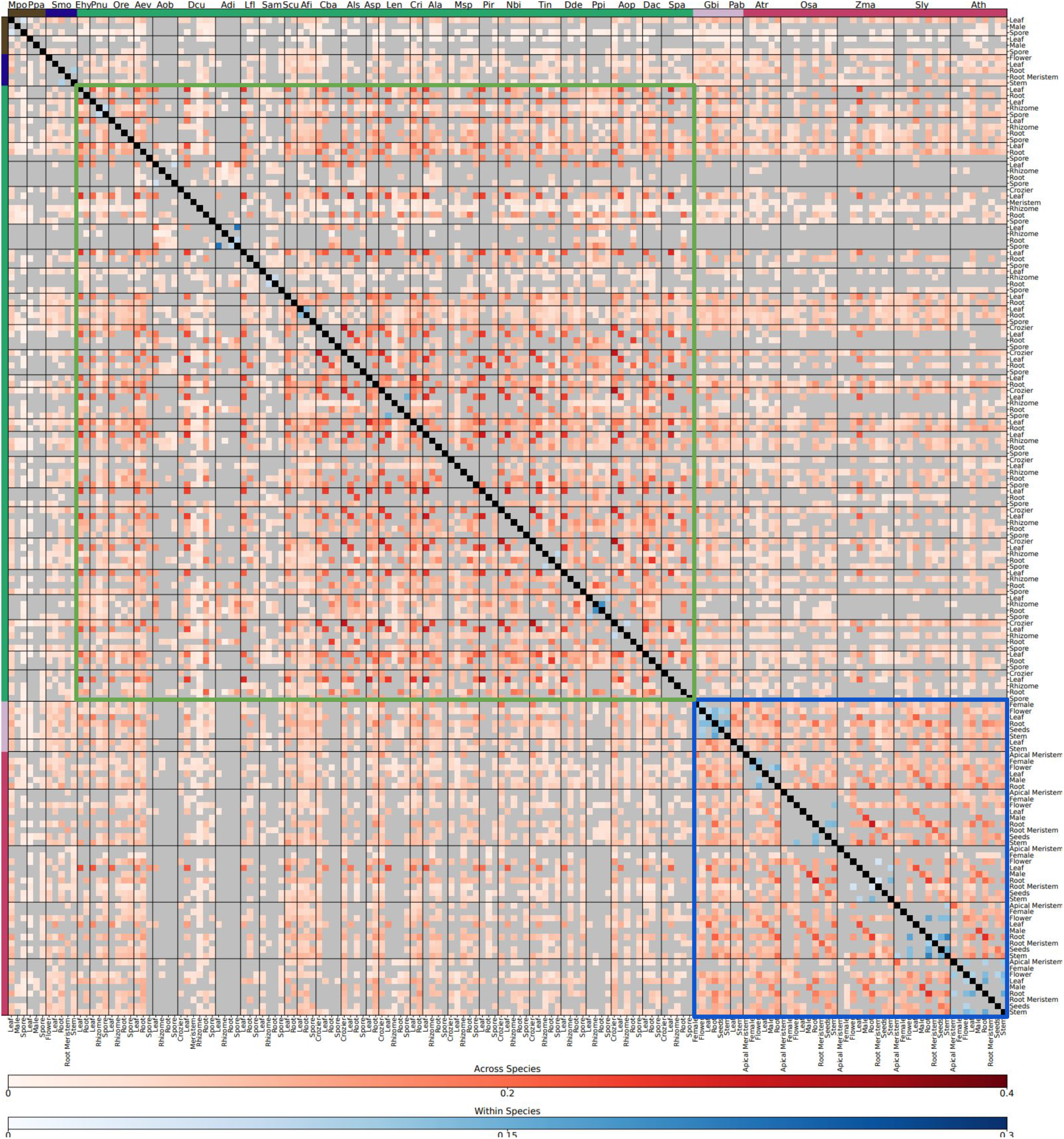
Transcriptome similarity comparison of Archaeplastida. The heatmap shows the conservation of organ-specific orthogroups across the species. The jaccard index of across species similarities are indicated by red shades, and for within species similarly with blue shades.

**Figure S9.**
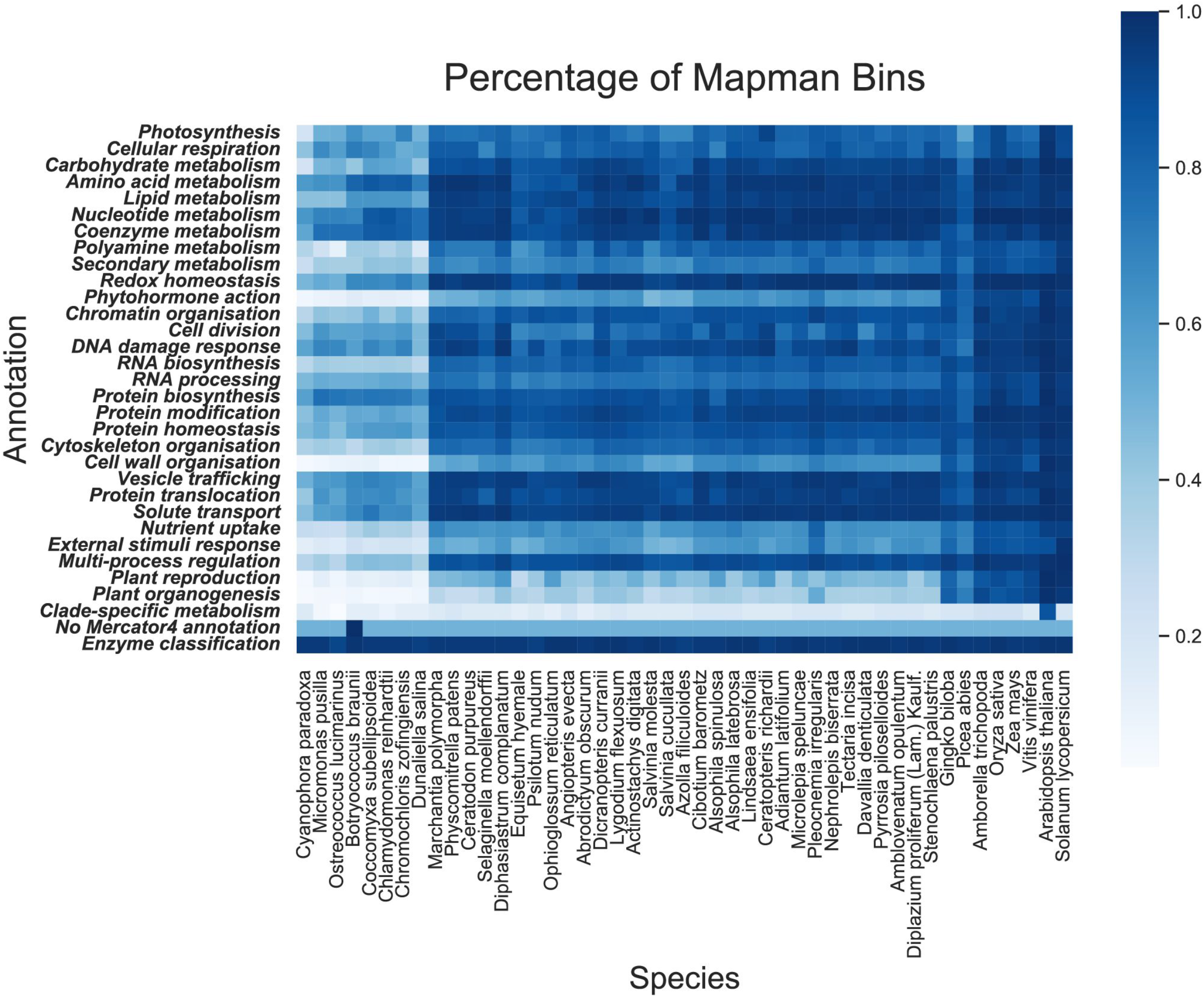
Gene functions found in Archaeplastida. Mapman bins (rows) found in the different species (columns). The colours indicate the fraction of found bins in a given species, where 1 indicates that all genes in a given bin are present, while 0 indicates complete absence.

**Figure S10.**
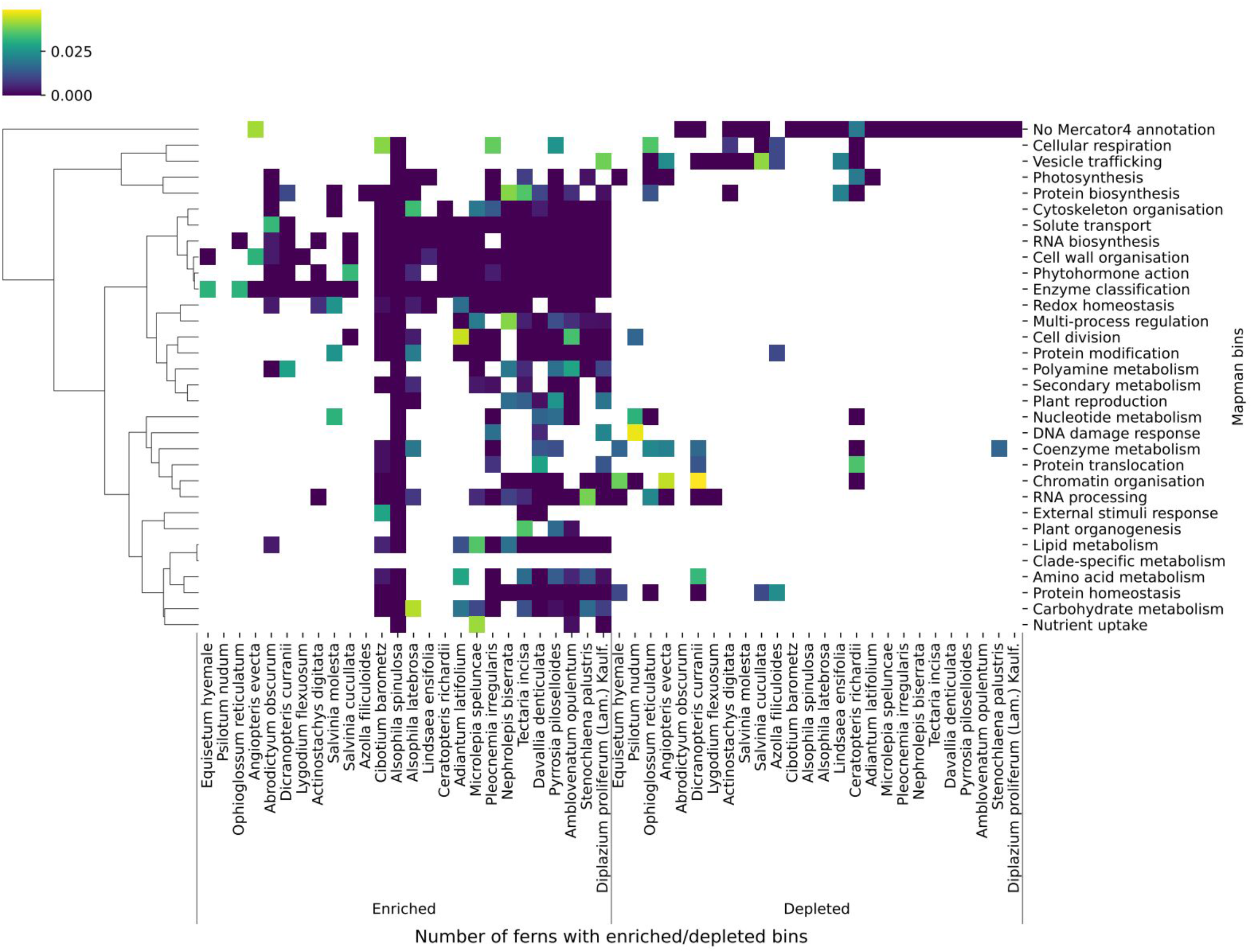
Enrichment and depletion of biological processes in neighbours of fern-specific genes. Clustermap showing significantly enriched and depleted primary Mapman bins (y-axis) in neighbours of fern-specific genes across ferns. The colour map indicates the significance of the biological processes, with yellow representing a p-value of 0.05 and blue representing a p-value of 0.00. P-values above 0.05 are masked.

**Figure S11.**
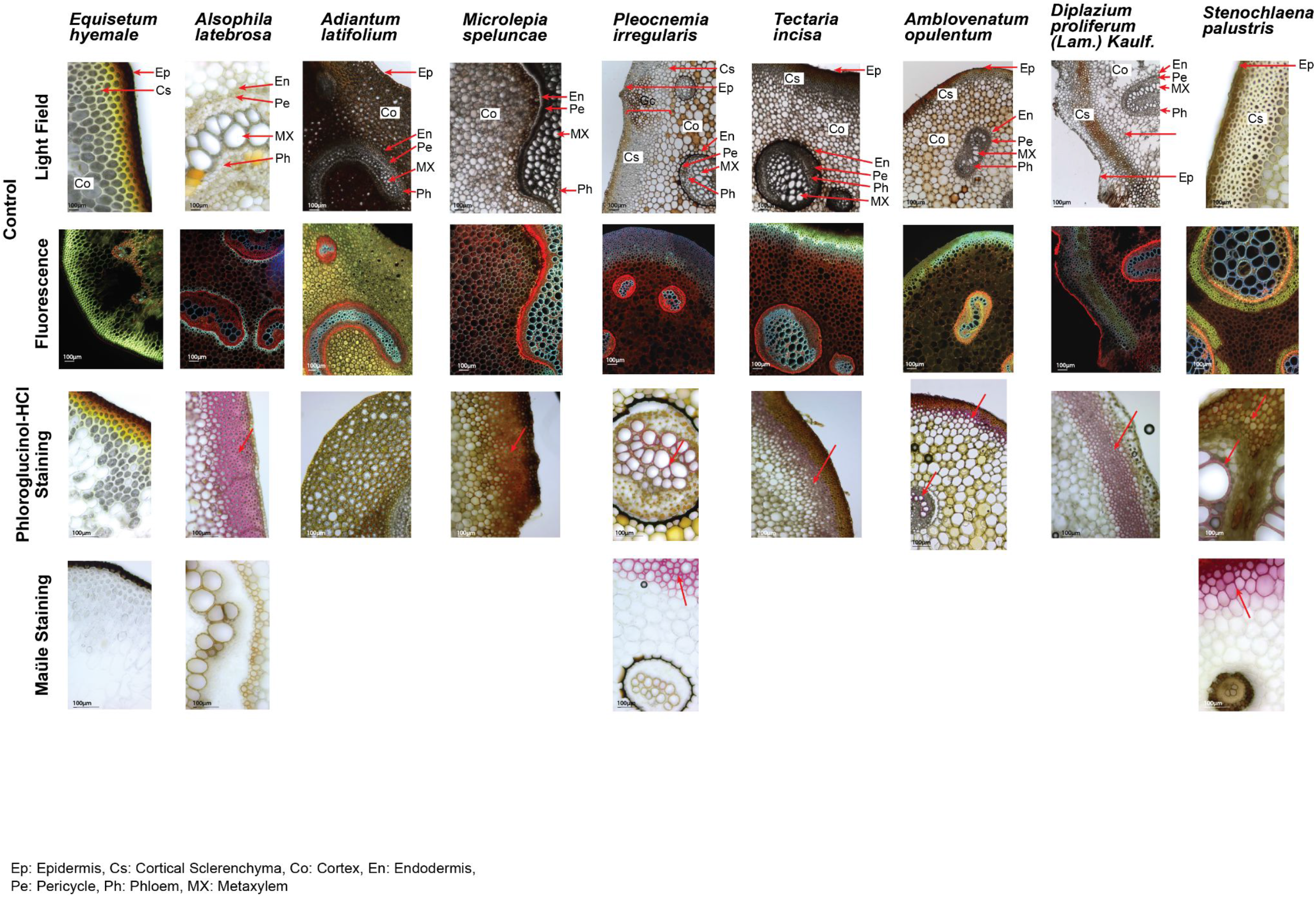
Light field, fluorescence, phloroglucinol and Maule staining of the selected ferns.

**Figure S12.**
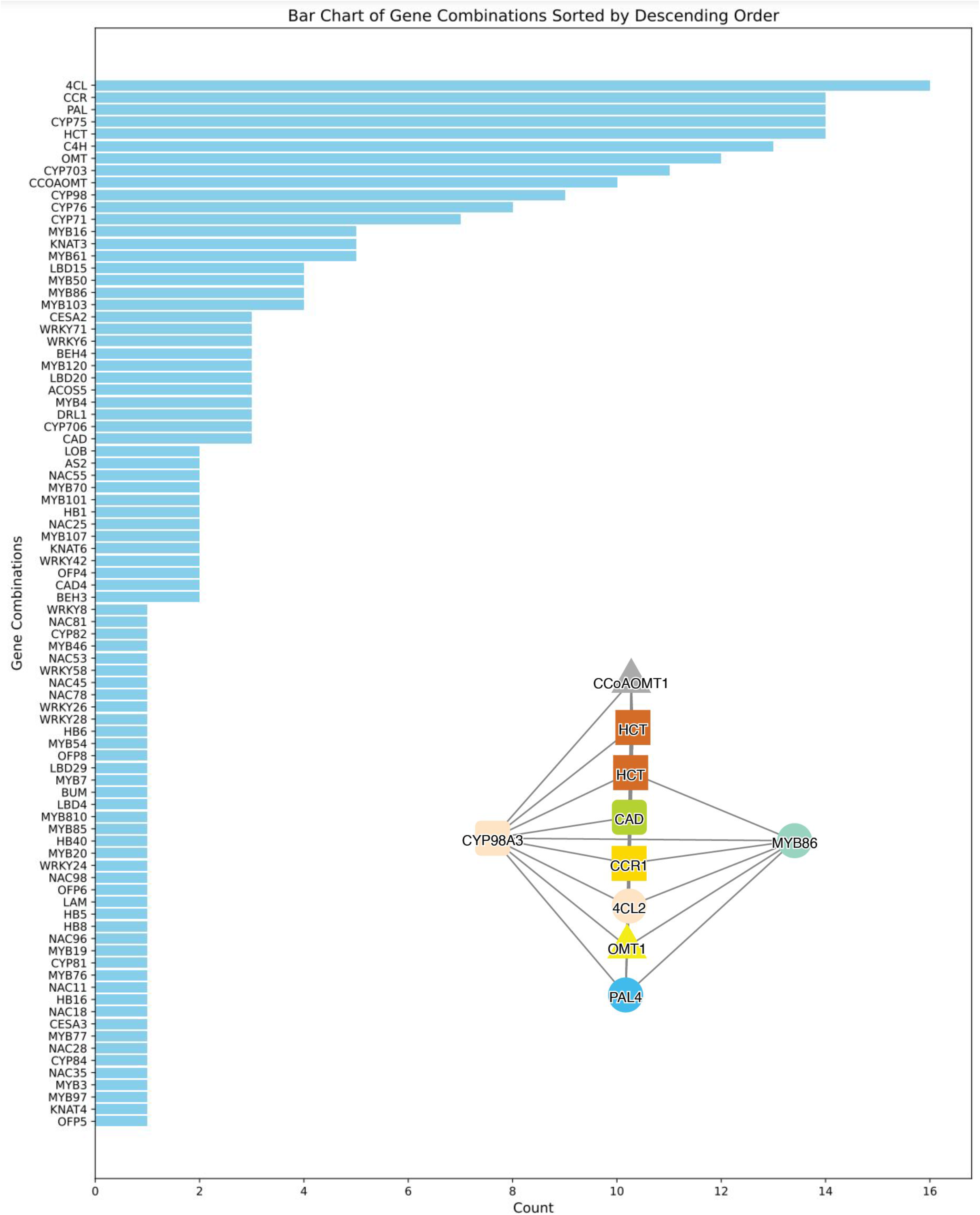
The number (x-axis) of the transcription factors, CYP450s enzyme families and lignin-related enzymes co-expressed with at least two lignin biosynthetic enzymes in the analyzed ferns. Co-expression network of PAL4 (blue circle) from Dicranopteris curranii. CYP96A3 and MYB86 are connected to seven and five enzymes involved in lignin biosynthesis.

**Figure S13.**
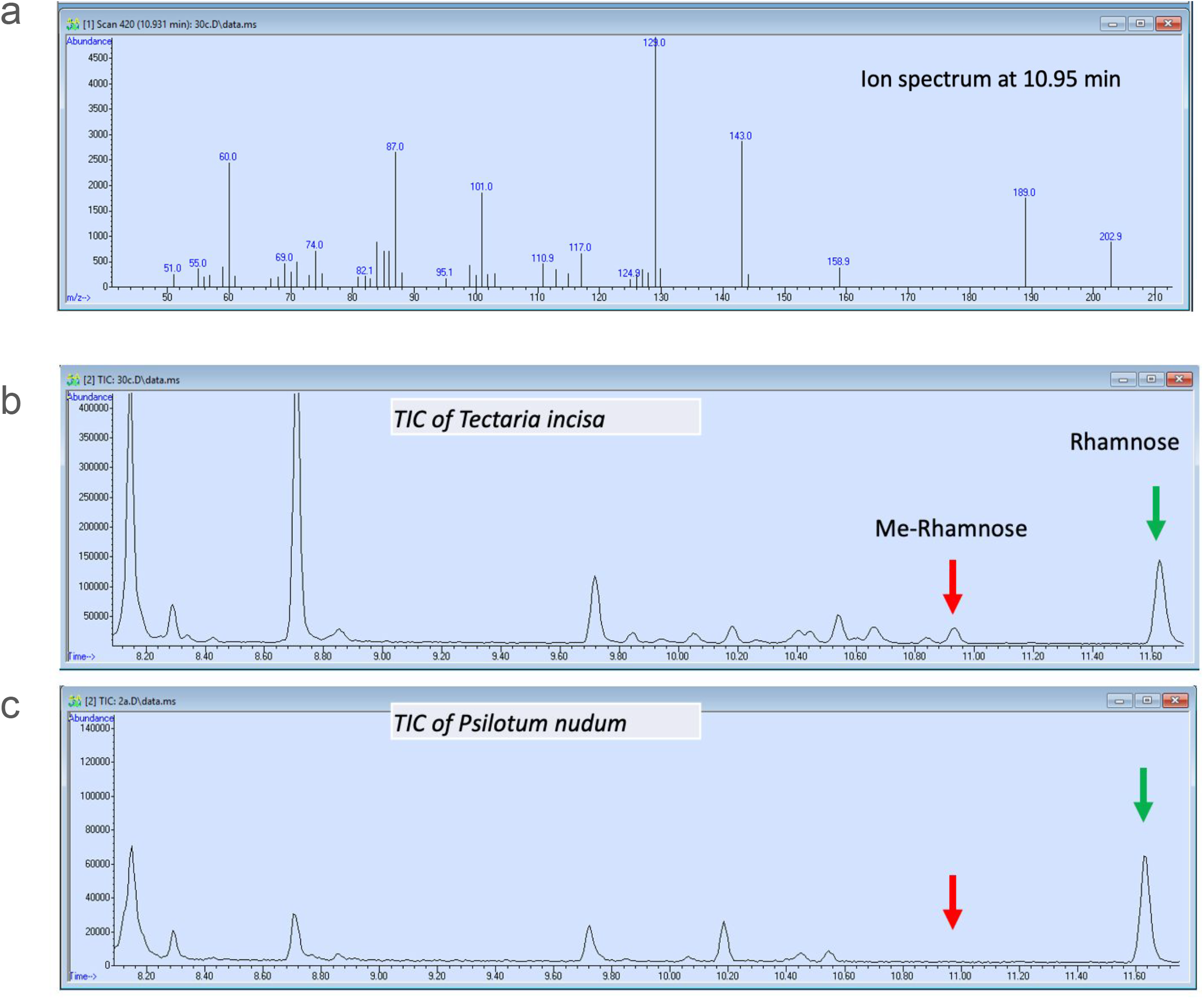
GC-MS analysis of 3*O*-MeRha*p*. a) MS profile of 3*O*-MeRha*p*. b) T*ectaria incisa* contains 3*O*-MeRha*p* (read arrow) and rhamnose. c) Psilotum nudum contains rhamnose but no 3*O*-MeRha*p*.

**Figure S14.**
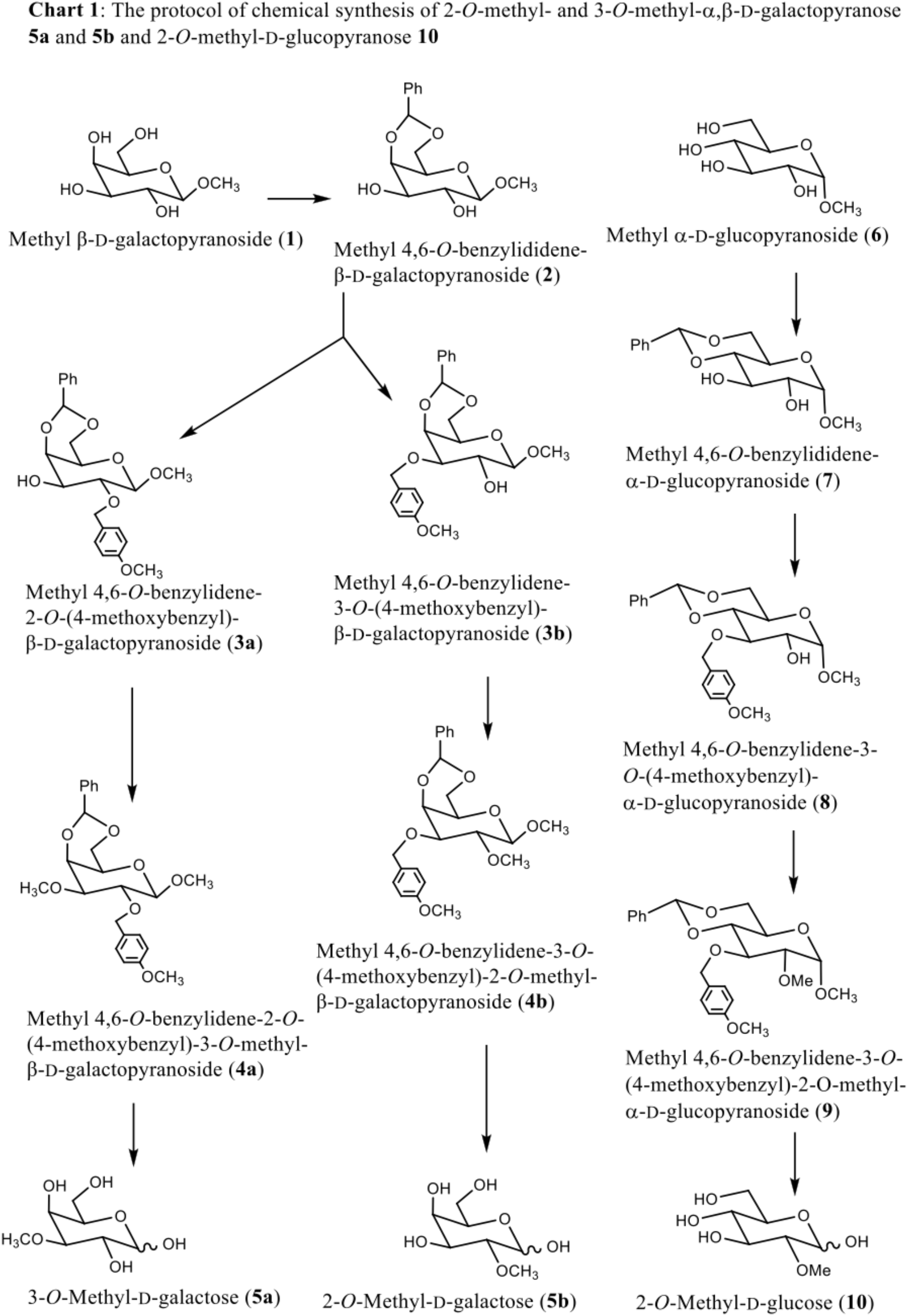
The protocol of chemical synthesis of 2-O-methyl- and 3-O-methyl-α, β-D-galactopyranose 5a and 5b and 2-O-methyl-D-glucopyranose 10.

**Figure S15.**
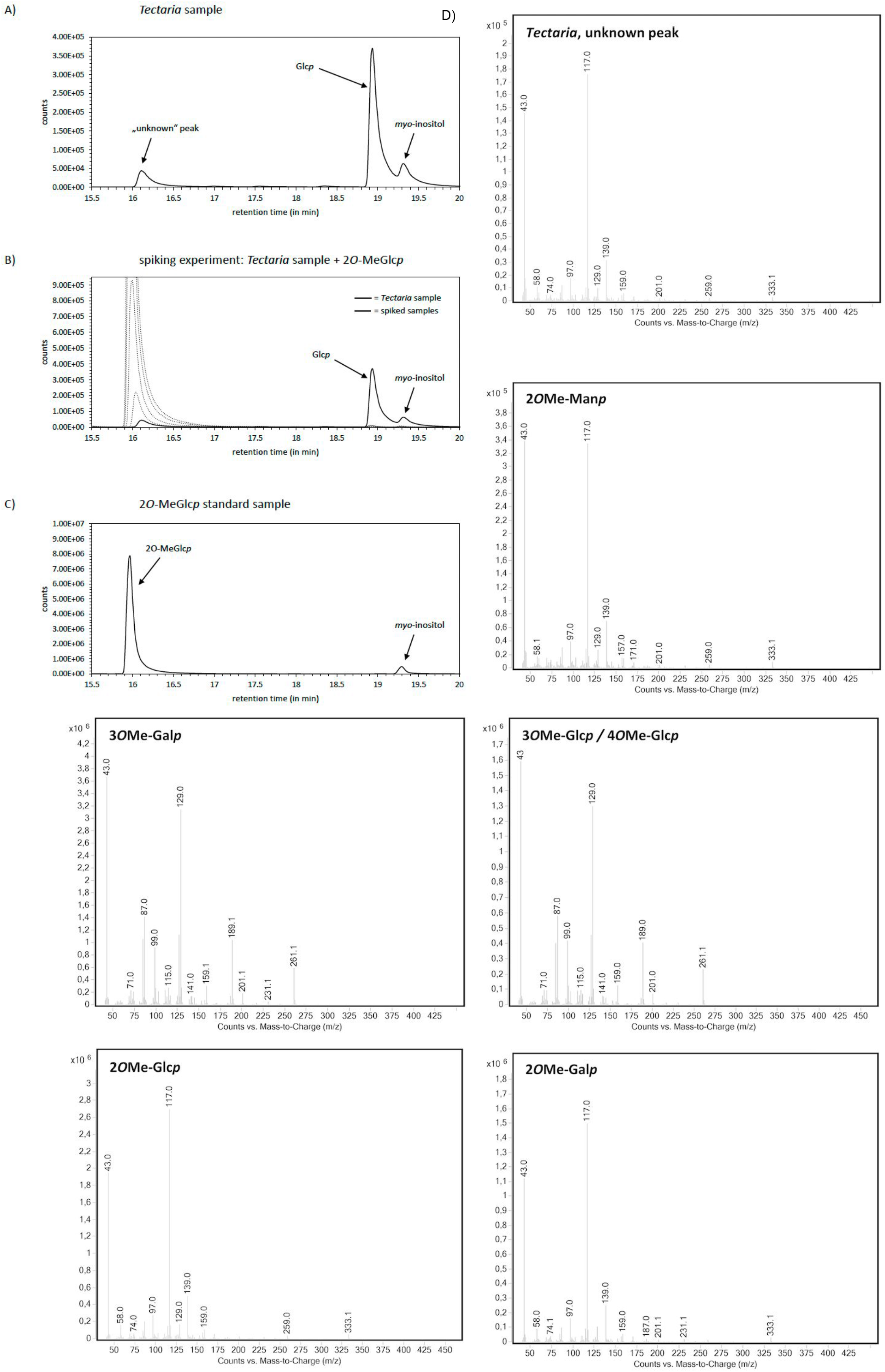
GC-MS analysis of cell wall sugars. a) Profile of T*ectaria incisa*, b) T*ectaria incisa* and 2*O*-MeGlc*p* standard and c) 2*O*-MeGlc*p* standard. d) GC-MS spectra of the unknown peak and the methylated sugar standards.

**Figure S16.**
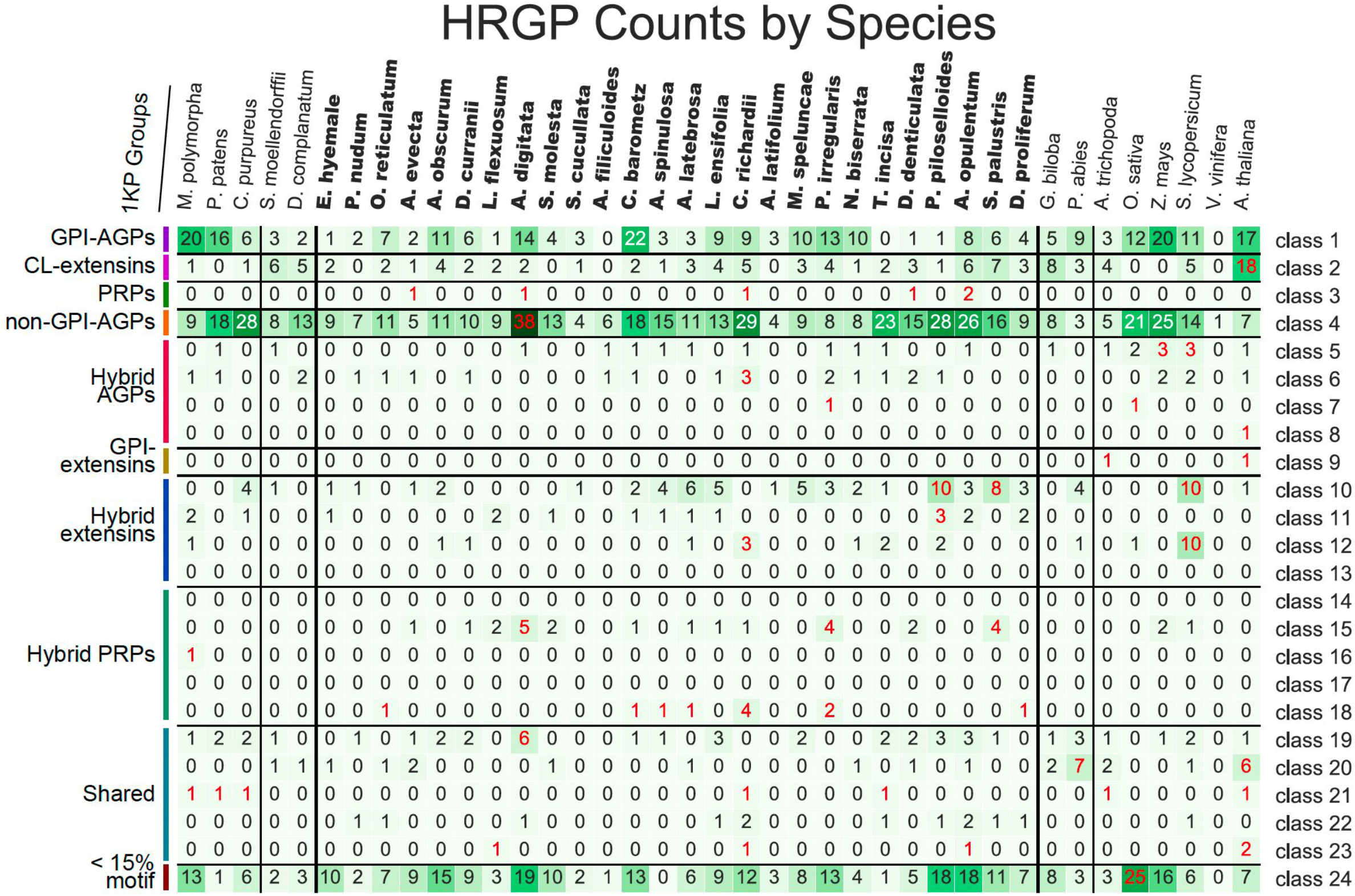
Gene copy number analysis of hydroxyproline-rich glycoproteins (HRGPs). Columns represent species, while rows correspond to a given class of HRGP. Red and blue numbers indicate that a given species contains significantly more/less genes than others.

**Figure S17.**
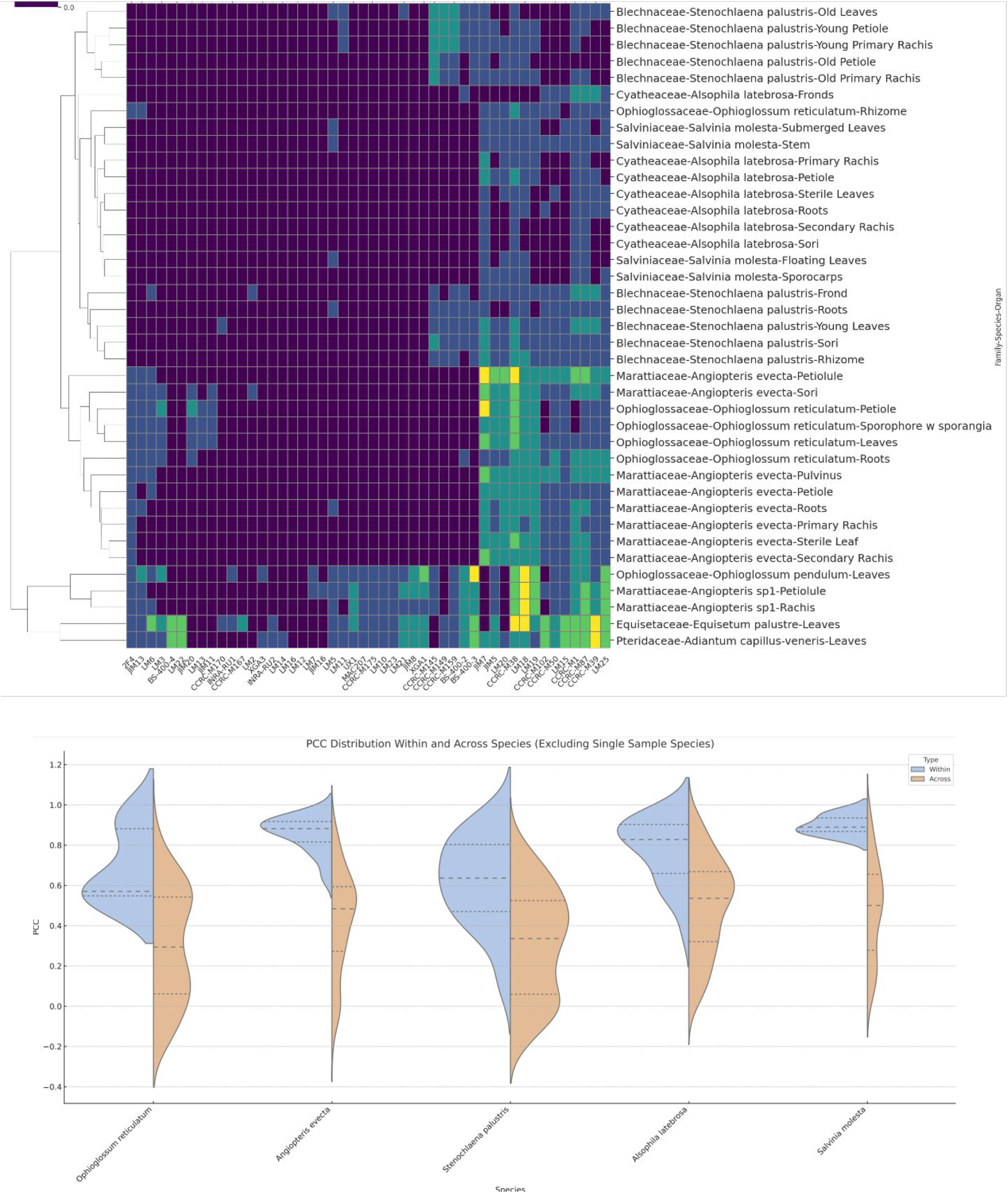
CoMPP analysis of ferns. a) The clustermap shows fern samples (rows) and antibodies (columns). The cells indicate the signal scaled from 0 (dark blue) to 1 (bright yellow). b) Pearson Correlation Coefficient (PCC) distribution of CoMPP profiles within (blue) and across (brown) species.

**Figure S18.**
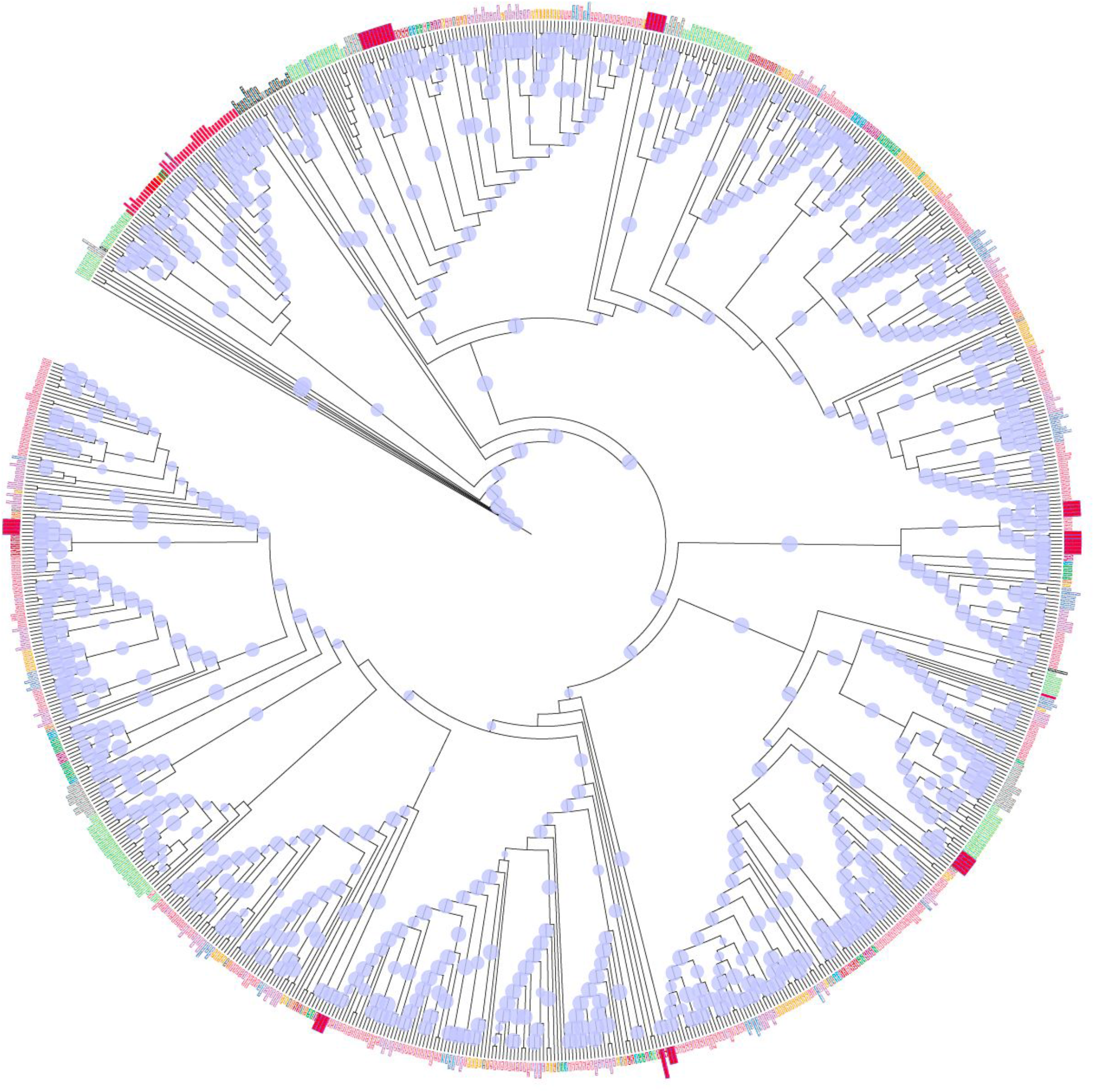
Phylogenetic analysis of CESA genes. The blue circles represent bootstrap values (value <50 are not indicated by a circle). The leaf colors represent the different species and orders.

**Figure S19.**
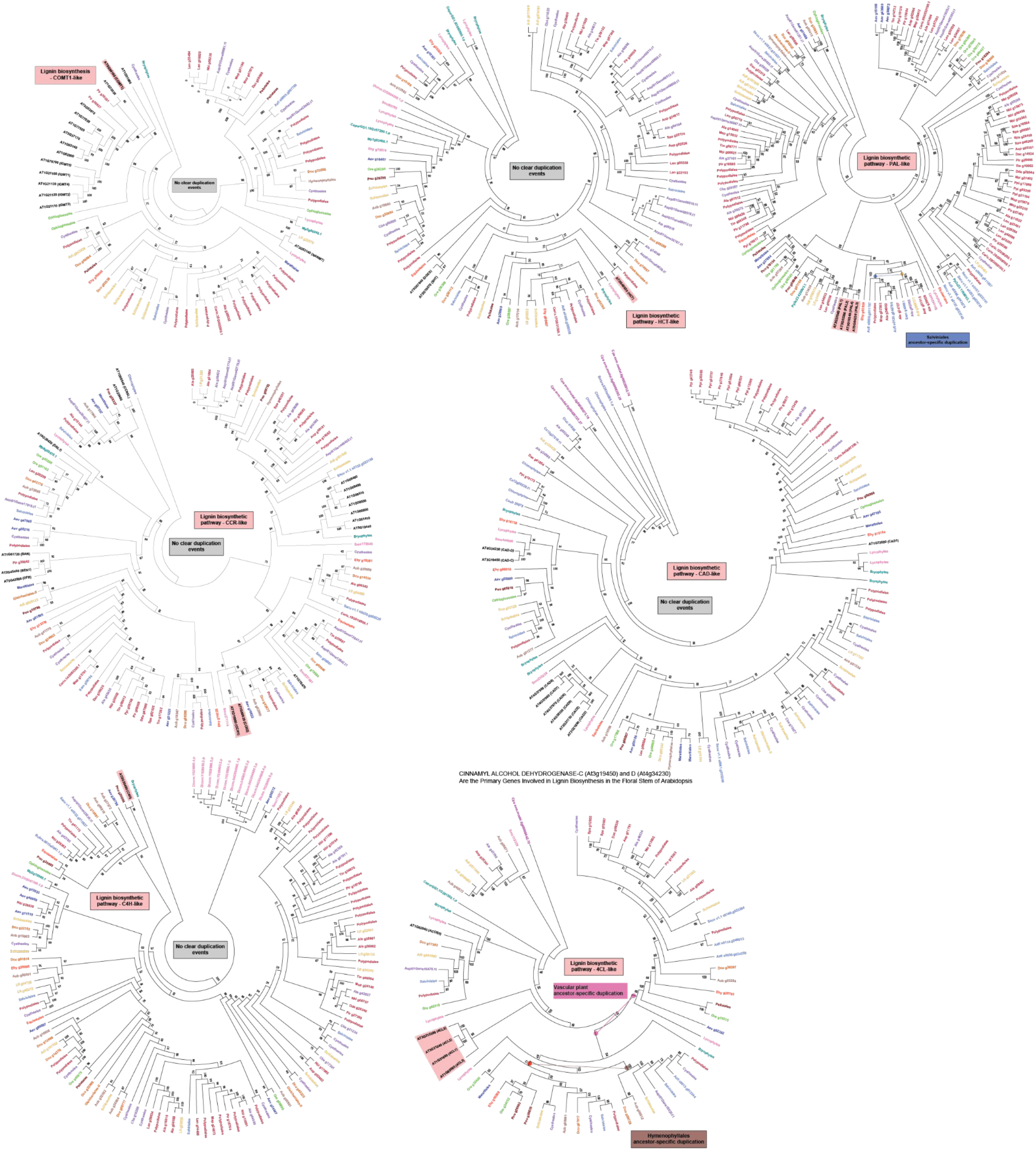
Phylogenetic analysis of lignin biosynthetic genes. Any inferred duplication events are indicated.

## Notes

https://conekt.plant.tools/

### Competing Interest Statement

The authors have declared no competing interest.

